# ‘Same same but different’ Exploring evolutionary and ecological causes of eye diversification in bumblebees

**DOI:** 10.1101/2022.10.25.513654

**Authors:** Pierre Tichit, Andrew Bodey, Christoph Rau, Emily Baird

**Affiliations:** Department of Biology, Lund University, Lund 223 62, Sweden; Diamond Light Source, Oxfordshire, United Kingdom; Department of Zoology, Stockholm University, 114 18 Stockholm, Sweden

**Keywords:** X-ray microtomography, bumblebees, compound eyes, vision, sensory ecology, eye evolution, phylogeny, functional traits, inter-specific variation, social parasitism, habitat drivers, sensory-based conservation

## Abstract

Bees rely heavily on vision during most of their interaction with the environment, but so far, visual abilities have not been included into functional investigations of these crucial pollinators. This is probably due to the lack of comprehensive and phylogenetically-controlled quantification of visual traits across species. In the present study, we used high-throughput micro-CT tools to quantify, compare and understand the diversity of visual traits of compound eyes in bumblebees. Visual systems of bumblebees were far from identical, with variations across sizes, castes and species. While phylogenetic proximity poorly supported interspecific variations, these were better explained by two ecological factors: social parasitism and habitat. The eye parameter – a metric that measures the relative investment of a compound eye into resolution or sensitivity – was lower in queens of social parasitic species than of non-parasitic species. Workers of species associated with forested habitat had distinct visual traits, including a higher eye parameter, than those of species living in open landscapes. These diverse visual traits are likely to provide selective advantages to bumblebees given their specific ecological requirements. We thus propose that social parasitism and forest habitat are drivers of the diversification of compound eyes in bumblebees. Finally, we discuss how the present study can inspire trait-based approaches in ecology and conservation biology.

## Introduction

To understand how living organisms interact with their environment across space and time, it is valuable to quantify the diversity of their generalizable functional properties, or traits. By investigating how traits of organisms are influenced by and influence the environment, functional ecologists achieve a broad and mechanistic understanding of the structure and functions of communities of organisms (Wong et al., 2019). Trait-based approaches can in turn generate predictions that inform conservation strategies to preserve endangered organisms and ecosystems (Chown, 2012; Kosman et al., 2019; Pacifici et al., 2020). For example, trait-based risk assessments investigate how traits determine the vulnerability of organisms to anthropogenic disturbances (e.g. landscape transformation, climate change, invasive species). In order to obtain comprehensive results, trait-based approaches should *(1)* integrate a wide range of traits that could play a role in the ecological question at hand. For terrestrial invertebrates, these traits are typically categorised in five groups: morphology, feeding, life history, physiology and behaviour (Moretti et al., 2017). Trait-based approaches must *(2)* perform comparisons across a sufficient number of organisms with some level of variability in the investigated traits. Finally, trait-based approaches should *(3)* be phylogenetically controlled, in order to disentangle whether similar functional properties likely result from shared environment or shared ancestry.

In animals, sensory systems gather information about the environment. This information is then processed by the brain and informs behaviours that in turn influence the type of information perceivable in the environment. For example, a bee perceives the odour of a flower, thus extends its tongue into the nectar spurs, an action that subsequently generates additional gustatory and mechanical cues. Because senses are the only interface between an animal and the environment, sensory systems have evolved in relation to specific ecological needs for producing the most adaptive behaviour whilst reducing the energetic costs incurred (Niven and Laughlin, 2008; Wehner, 1987). Despite their importance, sensory systems are overlooked in trait-based analyses. This is likely because sensory traits are often difficult to measure, and the available data on how they relate to an animal’s ecology are scarce. However, in recent years, sensory biologists have shown a growing interest for comparative approaches that quantify and integrate sensory properties into a phylogenetic framework (for example in arthropod eyes: Farnier et al., 2015; Feller et al., 2020; Keesey et al., 2020; Scales and Butler, 2016; Streinzer and Spaethe, 2014a). Comparative studies not only are valuable to understand the evolution of sensory systems (Chittka and Briscoe, 2001; Dangles et al., 2009), but also open the possibility to integrate sensory properties in trait-based analyses in ecology and conservation biology. This is what the present work starts to do by quantifying and integrating the variability of visual traits of bumblebees into an evolutionary and ecological framework.

There are about 250 species of bumblebee all within the genus *Bombus* (Goulson, 2010). In spite of being closely related, bumblebees live in different habitats, ranging from cluttered tropical forests (e.g. *Bombus transversalis*) to featureless tundra (e.g. *Bombus polaris*) (Goulson, 2010). Bumblebees are eusocial insects with three castes. Labour is typically divided between a queen that lays eggs and initially forages intensely to feed the first workers that, once they are in sufficient numbers (from a handful up to several hundreds), perform most of the tasks in the hive (Goulson, 2010). Males emerge at the end of the season and leave the hive to mate with newly emerged queens. There is an exception to this life-cycle in social parasitic – or cuckoo – bumblebees, where queens take over hives of their host species that will then rear their offspring (Lhomme and Hines, 2018). Bumblebees are large, hairy, tolerant to cold and adverse weather, and capable of buzz pollination, which makes them important pollinators, particularly in northern latitudes (Goulson 2010). Unfortunately, they have undergone an alarming decline over the past decades because of anthropogenic drivers such as landscape and climate change (Rasmont et al. 2015; Sirois-Delisle and Kerr 2018).

Like many flying insects, bumblebees rely heavily on vision for most behaviours. In flight and during landing, bumblebees use visual motion cues to control their position in space and their speed (Baird et al., 2011; Dyhr and Higgins, 2010; Linander et al., 2015; Reber et al., 2016). They use visual information to detect and distinguish flowers (Goulson, 2010), navigate between their hive and a food source (Mandal, 2018) and for mate-finding behaviour (Paxton, 2005). In addition to being visually-driven animals, bumblebees have a resolved phylogeny (Cameron et al., 2007), known behaviours, distinct habitats and a conservation importance, which makes them good candidates for comparative studies of visual systems.

To perform visually-guided behaviour, bumblebees possess two types of eyes: three ocelli and a pair of apposition compound eyes. The compound eye is composed of thousands of optical units called ommatidia. The ability of a portion of the eye to resolve details in a visual scene, or spatial resolution, is determined by the angle between two adjacent ommatidia, called the inter-ommatidial (IO) angle – such that a reduction in IO angle increases resolution (Land, 1997). The ability of a portion of the eye to see when light gets scarce, or light sensitivity, is in part determined by the diameter of the lens of the ommatidia, or facet diameter, such that an increase in facet diameter increases sensitivity. Other visual traits are meaningful to predict the visual abilities of bumblebees: for example, the facet number conditions resolution, whereas the extent of the field of view (FOV) indicates what portion of the world is visually sampled by the eye. The local optical properties of ommatidia vary across the topology of compound eyes and are optimised differently across the FOV in a way that often reflects the ecology of the animal (Cronin et al., 2014). To explore the relative investment in resolution and sensitivity, a useful metric is the local eye parameter – the product of facet diameter and IO angle. An eye parameter tending toward 0.29 μm.rad indicates a diffraction-limited region that maximises spatial resolution (Snyder, 1979). Conversely, an eye parameter above 1 μm.rad suggests a local investment towards improved sensitivity.

Using X-ray microtomography (micro-CT) – a high-throughput technique to generate fast and accurate 3D reconstructions of compound eyes (Baird and Taylor, 2017; Taylor et al., 2019) – we measure visual traits of compound eyes across an unprecedented number of bumblebees species. Despite the absence of ‘obvious differences in their visual system’ (Streinzer and Spaethe, 2014), we predict the presence of subtle interspecific variations of visual traits in bumblebees. We investigate whether the variability of bumblebees’ visual traits correlates with phylogenetic or ecological factors and examine the presence of evolutionary causality that support these relationships, which would suggest that phylogenetic or ecological correlates are drivers of eye diversification in bumblebees.

## Methods

### Sample collection

Most species and castes of bumblebees were collected from Scania province, southern Sweden, and around Abisko Scientific Research Station, northern Sweden. Specimens of *Bombus morio* were sampled from the Campus of the University of São Paulo, Brazil and specimens of *Bombus atratus* (also referred as *Bombus pauloensis*) kept in ethanol were obtained from a dataset published in Françoso et al. (Françoso et al., 2016). Workers of *Bombus transversalis* were caught at a field station of the national research institute of Amazonas, Brazil. To complete the dataset, dried bumblebees were obtained from the entomological collections of the biological museum at Lund University, Sweden. Finally, six workers of *Bombus terrestris* were specimens from commercial hives previously analysed in Taylor et al. (Taylor et al., 2019).

Specimens from outside the genus *Bombus* were queens of *Xylocopa tenuiscapa* collected and kept in ethanol near the Indian Institute of Science Education and Research Thiruvananthapuram, India, two *Apis mellifera* worker specimens previously analysed in Taylor et al. (Taylor et al., 2019) and three *Scaptotrigona depilis* worker specimens previously analysed in Tichit et al. (Tichit et al., 2020).

When living specimens were collected, they were promptly anesthetised with carbon dioxide for dissection.

### Sample preparation

For the freshly-collected specimens, the left compound eyes were dissected, fixated with a solution of paraformaldehyde and glutaraldehyde, stained with osmium tetroxide, dehydrated with a graded alcohol series, and embedded in epoxy resin as described in Taylor et al. (Taylor et al., 2019a). When the left compound eye was damaged, the right compound eye was prepared instead. Specimens stored in ethanol were directly dehydrated with a graded alcohol series followed by critical point drying. To allow subsequent registration of the eye in the world coordinates (see *volumetric and computational analysis*), bumblebee heads were dissected, fixated, dehydrated with a graded alcohol series and critical point dried.

### Sample scanning

All imaging using X-ray micro-CT was performed at the Diamond-Manchester Imaging Beamline I13-2 (Peić et al., 2013; Rau et al., 2011) at the Diamond Light Source, UK (beamtime numbers: MT13848, MT16052, MT17632-1, MT20385). Smaller bee samples (most compound eyes and the smallest heads) were imaged using x4 total magnification (providing a 1.6 μm effective resolution). Bigger samples (most heads and the largest eyes) were imaged with x2 total magnification (providing a 2.6 μm effective resolution). The scanning procedure was the same as in Taylor et al. (Taylor et al., 2019).

### Volumetric and computational analysis

The original images reconstructed from the scans were cropped and re-saved in 8-bit files in Drishti Paint (Limaye, 2012). All subsequent volumetric analyses were performed in Amira (FEI, Hillsboro, USA) where images were resampled at the voxel size of 4 μm or 5 μm. With a few exceptions, the procedure was generally identical to the method described in Taylor et al. (Taylor et al., 2019). In brief, this involved three main steps: labelling of eye and head features, measurement of facet dimensions, registration of the eye and head in world coordinates. The only difference with the previous method was that the inner layers of the compound eyes were not labelled, such that only the outer morphology of the eyes was exploited in the analysis. Instead, the eye volume was labelled semi-automatically using the segmenting tools in Amira. The outer surface of the cornea was then segmented from the eye label by manually drawing a geodesic path at the border of the cornea and extracting the enclosed surface. Similarly, the full head was also segmented in Amira.

The labels and measurements obtained during volumetric analysis were imported to *MATLAB* (The MathWorks Inc., USA) for computational analysis. Here again, the procedure follows the method described in Taylor et al. (Taylor et al., 2019) with two differences: *(1)* the mirroring of a compound eye (either because it was a right instead of a left compound eye, or to obtain the binocular overlap on the field of view of the left compound eye) was performed by flipping the eye with respect to the plane of symmetry of the head and *(2)* the validation of calculation was slightly less strict, as all calculations were repeated until the number of ommatidial axes was as close as possible to the facet number, but not necessarily always between 95% and 100% of the facet number.

For each individual, we used computational analysis to calculate the following global eye properties: the eye area (the surface area of the outer cornea of the eye), the number of facets, the extent of the field of view (FOV, defined as the percentage of the world sphere that the eye was sampling light from). Local eye properties that vary topologically over the FOV were also estimated: facet diameter, facet area, radius of curvature of the eye, corneal inter-ommatidial (IO) angle and eye parameter (the local product of IO angle and facet diameter). Local properties were mapped onto the world using a sinusoidal projection. Global values were calculated to summarise the local eye properties. These values included classical calculations such as the average or the median of each eye property across the surface of eye. To avoid distortions due to local outliers, minimum and maximum extremes were expressed as 10th and 90th percentiles of the data set and are referred to in the analysis as the lower and upper values of an eye property, respectively. All global values were referred to as visual traits in the analysis.

### Phylogenetical and ecological predictors

All subsequent analyses were performed in R (R Development Core Team, 2020) and all the code is reported in the supplementary material.

To perform a phylogenetic analysis of eye properties, we used a previously published high-resolution molecular phylogeny of bumblebees obtained from the analysis of nuclear and mitochondrial DNA sequences (Hines, 2008). The phylogeny was trimmed to the subset of bumblebee species investigated in this study. The two subspecies of *Bombus pascuorum smithianus* and *Bombus pascuorum pallidofacies* that were not present in the original phylogeny were included and separated by an arbitrary small distance. The phylogeny was converted into a covariance matrix that was used for Bayesian modelling (see *Data exploration and statistics*).

**Figure 1:**
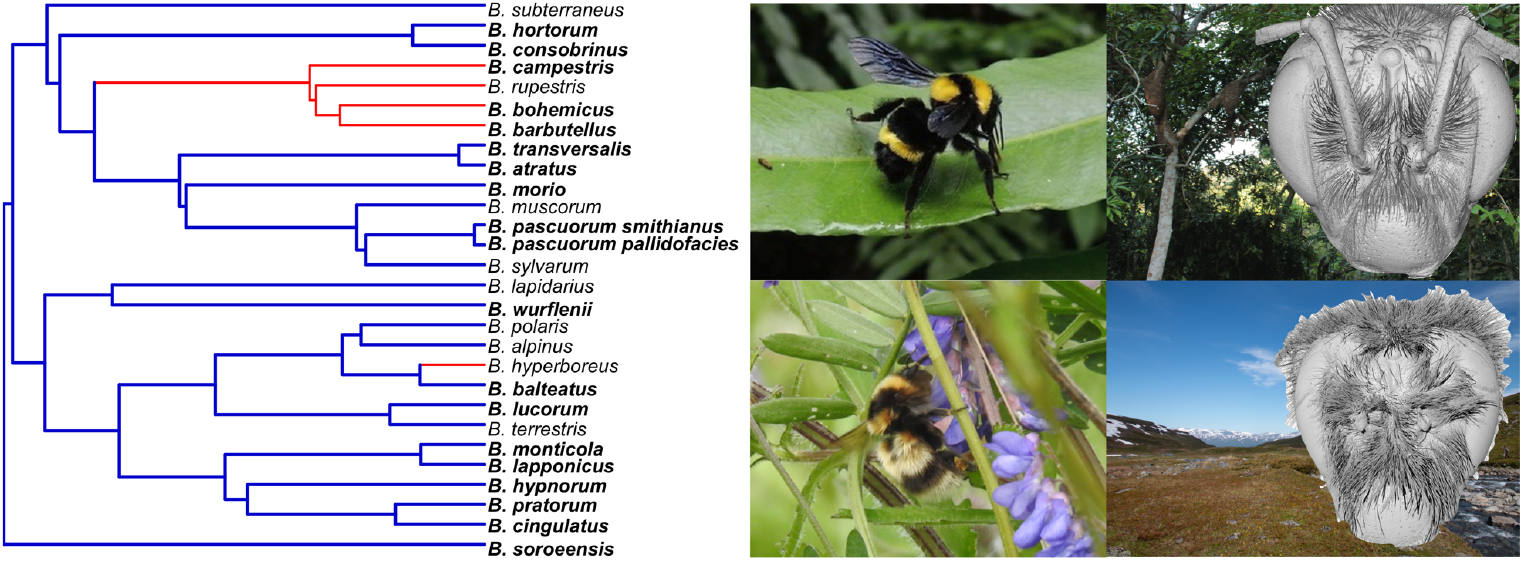
Phylogeny and ecology of bumblebees. *Left*: the phylogeny of 27 bumblebee species used in this study (Hines, 2008). Cuckoo bumblebees that perform social parasitism are marked in red, while species present in forest are in bold. Note that the two subspecies *Bombus pascuorum smithianus* and *Bombus pascuorum pallidofacies* that were not distinguished in the original phylogeny were separated by an arbitrary small distance. *Right:* example of the species *Bombus transversalis* (top, copyright John Asher / www.discoverlife.org) and *Bombus balteatus* (bottom) with their typical habitat and a volume rendering of their heads (grey images) obtained with micro-CT.

Three ecological predictors were collected: the presence of social parasitism (hereby referred as *parasitism*), the presence of the species in forested habitats, the affinity of the species with the forested habitat. The second variable, referred as *presence in forest*, was a binary variable scoring the association of a species with forested environments (regardless of its strength) as reported in the naturalist book ‘The bumblebees of Sweden’ (Mossberg and Cederberg, 2012). For the few non-Swedish species, we used the presence of forest in the vicinity of the capture sites to estimate the index. The third variable, or *forest affinity index*, was a continuous variable (scoring from 0 – never found in forest – to 1 – only found in forest) based on a published review of bumblebee habitats (Liczner and Colla, 2019). The index was calculated with the following equation:

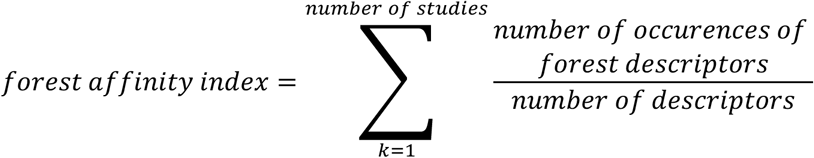

For example, if two studies were reviewed for a species, and the respective descriptors were “agricultural landscape, field margins, forest edges” and “gardens, forest”, the *forest affinity index* was 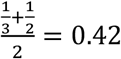. Forest descriptors could be “tropical forest”, “forest”, “woodlot”, “forest edge” and “forest interspace”. When data were not available in the review, we applied the same calculations to the habitat description from Mossberg and Cederberg (Mossberg and Cederberg, 2012).

### Data exploration and statistics

Standard statistical analyses – including Students t-tests, Shapiro-Wilk tests and Pearson’s product-moment correlation tests – were used to explore the variability of bumblebee eye properties. The coefficient of variation *CV_i_* and the half range *HR_i_* of variation in an eye property *i* were calculated with the following formulas:

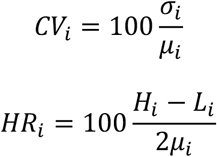

Where *μ_i_*, *σ_i_*, *H_i_*, *L_i_* are the mean, standard deviation, upper and lower values of the eye property *i* respectively. Allometric curves and estimates were obtained by fitting a standardised major axis to the log-transformed eye size and eye property *i* using the *SMATR* package (Warton et al., 2012).

All linear models in part II were performed using the Bayesian Markov Chain Monte Carlo Generalised Linear Mixed Models (*MCMC-glmm*) package (Hadfield and Nakagawa, 2010). All continuous response and explanatory variables were scaled and centred around zero prior to modelling. The phylogenetic covariance matrix and replication were the two random effects in the model. All scaled eye properties were sampled from a Gaussian distribution with uninformative priors. When modelled as response variables, the ecological predictors *parasitism* and *presence in forest* were sampled from a categorical distribution with the prior advocated in a previous publication (Villemereuil et al., 2012). *MCMC* simulations were run for 1 100 000 iterations with a burn-in of 100 000 iterations. Plots of the traces of the sampled output and density estimates were used to verify that there was little autocorrelation between successive iterations. The quality and significance of the models were assessed using Bayesian p-values (*p_MCMC_*) and the Deviance Information Criterion (*DIC*). The model with the lowest *DIC* was considered the best. In case of equality, the simplest model was favoured. The phylogenetic heritability was the proportion of the posterior covariance explained by phylogeny.

## Results and discussion

### Exploring the variability of visual traits in bumblebees

#### Quantification of the variability of visual traits in bumblebees

Our first goal was to assess the variability of visual traits across bumblebee species. To do so, we generated three dimensional optical models of compound eyes and used them to estimate visual traits for each of the 72 individuals across 27 species, including female workers (N = 37), female queens (N = 16) and males (N = 19) *(**Table S1**)*. On average, bumblebee compound eyes consisted of 5598 ommatidia with a facet diameter of 24 μm *(**Table 1**)*. The average lower, mean, and upper IO angles were 1.2°, 1.7° and 2.3° respectively. The local product of IO angle and facet diameter – called the eye parameter – indicates how compound eyes trade-off resolution and sensitivity. In this study, the average eye parameter across all the bumblebees was 0.71 μm.rad, indicating an overall investment towards relatively high resolution but low sensitivity consistent with the optical requirements of a fast-moving diurnal insect (Warrant et al., 2004).

**Table 1:** Summary of the visual traits of the bumblebees in the study (N = 72).

|  | **N** | **mean** | **Standard deviation** | **minimum** | **10% percentile** | **median** | **90% percentile** | **maximum** | **Coefficient of variation (CV)** | **Relative half range (HR)** |
| --- | --- | --- | --- | --- | --- | --- | --- | --- | --- | --- |
| **facet diameter ($\mu\text{m}$)** | 72 | 24.22 | 2.07 | 18.84 | 22.10 | 23.92 | 27.04 | 28.97 | 8.54 | 20.92 |
| **facet area ($\mu\text{m}^2$)** | 72 | 514.06 | 88.19 | 307.56 | 425.33 | 498.22 | 632.41 | 730.72 | 17.16 | 41.16 |
| **curvature ($\mu\text{m}$)** | 72 | 881.89 | 153.97 | 620.94 | 707.51 | 857.79 | 1115.81 | 1278.23 | 17.46 | 37.27 |
| **IO angle ($^\circ$)** | 72 | 1.69 | 0.19 | 1.20 | 1.51 | 1.68 | 1.93 | 2.13 | 10.94 | 27.56 |
| **facet number** | 72 | 5598 | 972 | 3916 | 4557 | 5418 | 6884 | 7667 | 17.36 | 33.50 |
| **eye parameter ($\mu\text{m}.\text{rad}$)** | 72 | 0.71 | 0.07 | 0.54 | 0.62 | 0.71 | 0.79 | 0.88 | 9.68 | 23.86 |
| **extent of the FOV (%)** | 72 | 23.79 | 2.71 | 18.51 | 19.99 | 23.90 | 27.33 | 28.95 | 11.39 | 21.93 |
| **lowest eye parameter ($\mu\text{m}.\text{rad}$)** | 72 | 0.47 | 0.04 | 0.32 | 0.43 | 0.47 | 0.53 | 0.57 | 9.40 | 26.16 |
| **highest eye parameter ($\mu\text{m}.\text{rad}$)** | 72 | 0.98 | 0.11 | 0.71 | 0.85 | 0.96 | 1.11 | 1.35 | 11.73 | 32.99 |
| **highest facet diameter ($\mu\text{m}$)** | 72 | 26.67 | 2.38 | 20.84 | 24.10 | 26.51 | 29.56 | 32.17 | 8.91 | 21.24 |
| **lowest IO angle ($^\circ$)** | 72 | 1.22 | 0.13 | 0.93 | 1.05 | 1.21 | 1.37 | 1.53 | 11.01 | 24.70 |
| **highest IO angle ($^\circ$)** | 72 | 2.25 | 0.29 | 1.49 | 1.94 | 2.25 | 2.61 | 3.14 | 12.81 | 36.51 |
| **Eye size ($\mu\text{m}$)** | 72 | 1692.99 | 273.02 | 1218.75 | 1392.29 | 1632.76 | 2085.72 | 2326.98 | 16.13 | 32.73 |

Are some visual traits more variable than others across bumblebees? To quantify the spread of each visual trait *i*, we calculated the coefficient of variation *CV_i_* and the half range *HR_i_*. Although *CV* and *HR* were slightly higher for curvature, facet number and eye size, there was little variation of these two metrics across visual traits (of the same dimension) *(**Table 1**)*. This indicates that most visual traits are equally variable across bumblebees.

To reduce the number of visual traits to be considered in further analysis, we computed a Principal Component Analysis (PCA) across all visual traits *(**Figure 2**)*. Most of the variability in the data (86%) was explained by the two first components the PCA. Unsurprisingly, some traits were highly correlated and formed five groups of covariates *(**Figure 2a-c**)*. In further analysis, we thus focused on the following variables: eye size, IO angle, facet number, eye parameter and facet diameter.

**Figure 2:**
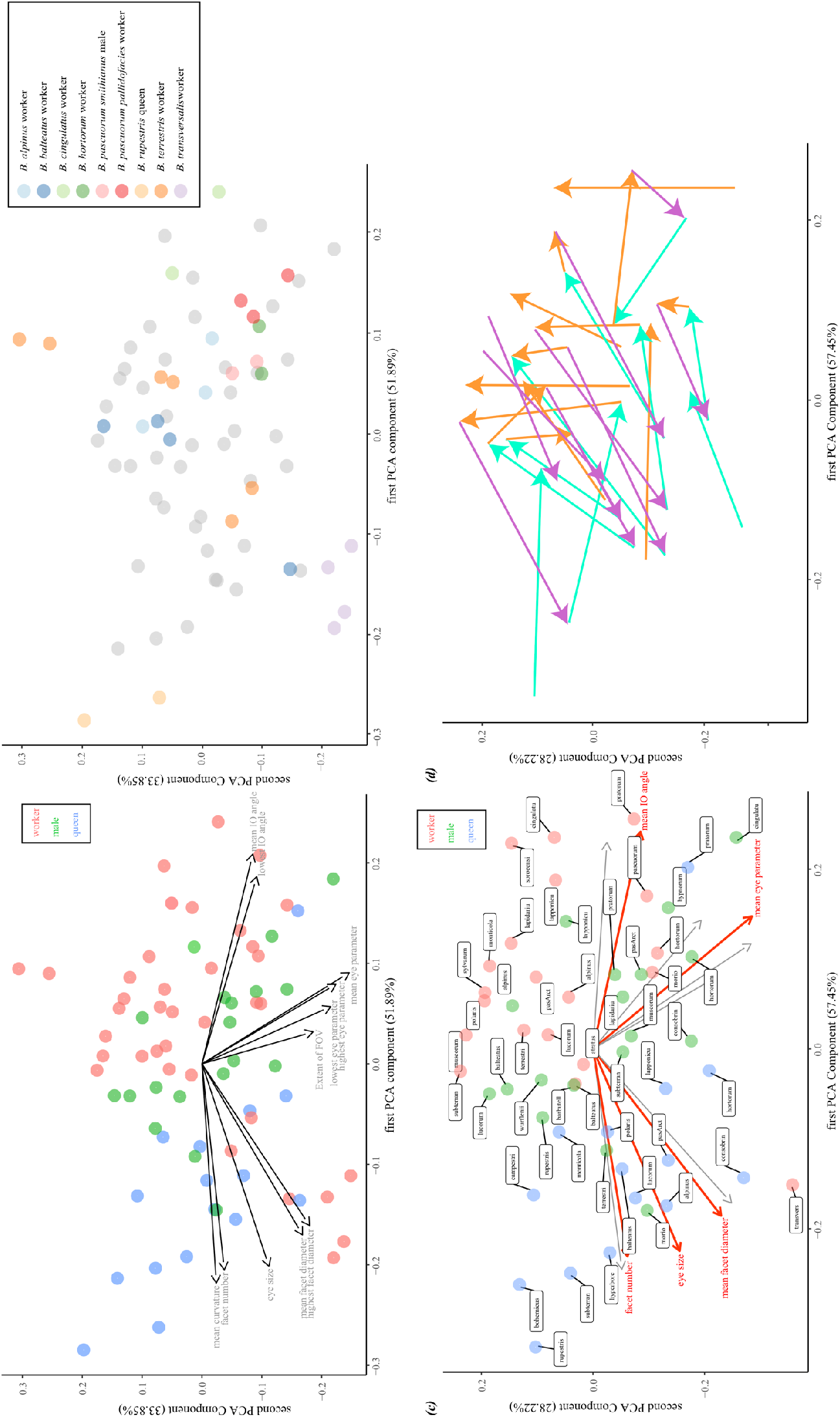
Exploring the variability of visual traits in bumblebees. Plots of the two first components of a Principal Component Analysis (PCA) over all visual traits. *(a-b)* Each point represents an individual bumblebee colour coded by caste *(a)* or replication category *(b)*. *(c-d)* the PCA was performed across categories, such that a point represents a species and caste of bumblebee *(c)*. In *(a)* and *(c)*, the arrows indicate the loadings of the visual traits. *(c)* This enabled to select only five variables for further analysis (red arrows). Note that the species label for *Bombus pascuorum smithianus* is ‘pasArct’, while ‘pascuorum’ refers to the subspecies *Bombus pascuorum pallidofacies. (d)* points (same as in *(c)*) were connected by species when possible: queen to male (cyan), male to worker (orange), worker to queen (purple). This revealed some species-specific similarities, in particular of workers-queens.

There were several categories of bumblebees in our analysis where more than one (and up to 6) replicates were available. Unsurprisingly, the average dissimilarity of visual traits (measured as the Euclidean distance in PCA space, ***Figure 2b**)* was lower between replicates than between non-replicate specimens (one-sided paired t-test, t = -4.4, df= 8, p < 10^-2^). As we expect, replicates of the same category to be most similar to each other, this finding confirms that our method accurately estimates visual properties (as was also clear from the three replicates of *Scaptotrigona depilis, **Figure 3**)*.

**Figure 3:**
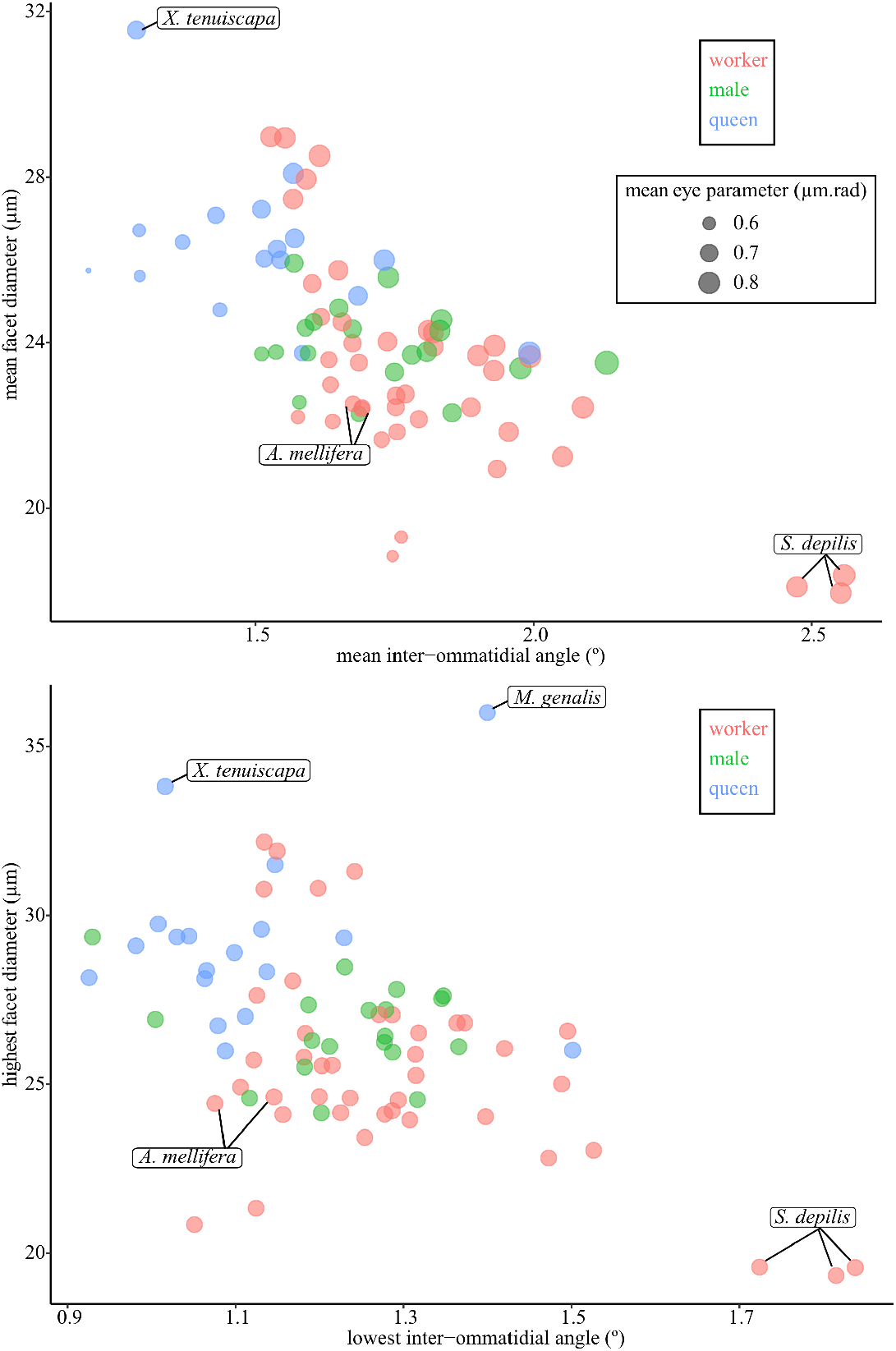
Comparing the variability of visual traits in bumblebees. The relation between facet diameter and inter-ommatidial angle in bumblebees (N = 72) can be compared to other bees: *Apis mellifera, Scaptotrigona depilis, Xylocopa tenuiscapa, Megalopta genalis*. In the upper graph, the size of each point represents the eye parameter.

#### Comparison to previous studies on compound eyes

How do the visual traits measured in this study compare to previous work on bumblebees? The average facet diameters and numbers obtained here are similar to the ones found in males and workers of *Bombus terrestris* (Kapustjanskij et al., 2007; Spaethe, 2003; Taylor et al., 2019). We also compared the values of eye area, mean facet diameter and facet number for seven species obtained here and analysed in a previous study (Streinzer and Spaethe, 2014). While the facet number was identical between this study and Streinzer and Spaethe (paired t-test, t = 0.66, df = 12, p = 0.52), eye area was marginally higher (paired t-test, mean difference = 0.23 mm^2^, t = 2.29, df = 12, p = 0.040) and mean facet diameter was higher (paired t-test, mean difference = 4.4 μm, t = 14.9, df = 12, p < 10^-8^) in Streinzer and Spaethe, with the variation likely being due to methodological differences. The lower, mean, and upper IO angles were in the range of the extrema (0.6° to 3.3°) measured previously with the pseudo-pupil method in workers of *Bombus terrestris* (Spaethe, 2003), which is consistent with our estimates. The overall agreement between the visual trait measurements obtained here and those of previous studies using different approaches further suggests that our method does indeed provide reliable results.

How do bumblebee visual traits compare to those from bees in other bee genera? To answer this, we analysed three *non-Bombus* species from the family Apidae *(**Table S2**)*, and the Halictidae *Megalopta genalis* (using measurements obtained from (Warrant et al., 2004)). The eye properties of the honeybee *Apis mellifera* were similar to bumblebees of equivalent size (facet diameter, corneal IO angle and eye parameter *(**Figure 3**)*, whereas the large carpenter bee *Xylocopa tenuiscapa* and the small stingless bee *Scaptotrigona depilis* were characterized by higher (respectively, lower) facet diameters and lower (respectively, higher) IO angles. Despite lying outside of the range observed for bumblebees, these two diurnal bees had a mean eye parameter similar to bumblebees (between 0.7 μm.rad and 0.8 μm.rad, ***Figure 3**, **Table 1**)*, indicating a similar investment towards relatively high resolution and low sensitivity. On the contrary, the crepuscular bee *Megalopta genalis* possesses higher facet diameter and IO angles, clearly lying outside of the relationship observed in diurnal bees *(**Figure 3b**)*. This trend is reflected by an enhanced upper eye parameter (1.2 μm.rad), indicating an investment towards improved sensitivity at the cost of resolution. Taken together, these comparisons with other bee genera indicate that the range of visual traits observed in bumblebees matches with the broad ecological requirements of an insect flying in daylight.

How do bumblebee visual traits compare to those of species in other insect orders? Using previous literature, we calculated the variability of visual traits *(CV* and *HR)* in a genus of damselflies (Scales and Butler, 2016) and a family of psyllids (Farnier et al., 2015). We found that bumblebee visual traits were approximately equally variable *(**Table 1**)* as in psyllids *(**Table S3a***, only *HR* and not *CV* is relevant given the small number of studied psyllids) and in damselflies *(**Table S3b**)*. In these damselflies and psyllids, the variability of visual traits was probably partly driven by the association of species with specific micro-habitats. Whether habitat is also a driver of the equally variable visual traits in bumblebee is unknown, and this possibility is explored in more detail in II). Regardless of what causes this variability, our study indicates that – despite being restricted in one genus and seemingly similar – bumblebee compound eyes are far from all identical.

#### Influence of size, caste and species on visual traits in bumblebees

IO angle, facet number, and facet diameter were significantly affected by eye size *(**Table S4*** and ***figures S2**)*. Interestingly, the eye parameter seemed poorly explained by eye size (p = 0.089), suggesting that the variability of this trait may correlate with other factors. The scaling exponents for facet number (1.09) and upper facet diameter (0.56) in this dataset were close to the scaling exponents for facet number (1.39) and maximum facet diameter (0.26) calculated from Streinzer and Spaethe (Streinzer and Spaethe, 2014; Taylor et al., 2019). These allometric effects underline the necessity to include eye size when investigating drivers of visual traits.

Our analysis confirms that bumblebee visual traits are influenced by caste *(**Figure 2c**)*. Workers (with the exception of *Bombus transversalis*) had the lowest values for eye size, facet number and diameter, queens (except in *B. pratorum)* had the highest values for eye size, facet number and diameter, and males had intermediate values. These relationships are likely due to the close allometric association between eye size, facet number and diameter described earlier *(**Table S4*** and ***figures S2**)* and are consistent with previous comparisons in bumblebees (Streinzer and Spaethe, 2014). The differences in visual traits between castes also motivates the need to treat them separately in the following analysis.

Interestingly, the variations of eye parameter (and, to a lesser extent, of IO angle) seem better explained by species than by caste, as illustrated by connecting queens, males and workers of the same species in PCA space *(**Figure 2d**)*. In other words, individuals of the same species but two different castes seemed more similar than individuals of other species regardless of their caste (one-sided t-test on dissimilarity in PCA space: males vs workers, t = -3.5, df = 38, p < 10^-3^, males vs queens, t = -1.6, df = 38, p = 0.056, workers vs queens, t = -1.7, df = 31, p = 0.051). This suggests the presence of species-specific drivers of visual traits and, in particular, of eye parameter and IO angle. One possible driver is phylogenetic history, i.e. that closely related bumblebee species share more similar eye properties than distant species due to phylogenetic inertia. Another possible driver is ecology because species sharing similar visual ecologies likely have visual traits that are tuned to similar selective pressures.

### Phylogenetic and ecological drivers of visual traits in bumblebees

#### Link between phylogeny and visual traits

To quantify how much of the variability of bumblebee visual traits can be explained by phylogenetic history, we fitted a Bayesian linear model to each property that solely included replication and a molecular phylogeny as two random effects (null model). We found that the typical phylogenetic heritability is low, (<1%, ***Table S5**)* and that the credible intervals were typically large, with most upper bounds above 90%. This indicates that the effect of phylogeny on visual traits is probably low and a large part of the variability remains to be explained by factors other than phylogenetic relationships, although the effect of phylogeny is too uncertain to be neglected in further analyses.

#### Possible ecological drivers of visual traits

We explored whether ecological differences between bumblebee species explain the variability of visual traits. We hypothesised that the compound eyes of bumblebees with social *parasitism* (all members of the *Psithyrus* subgenus and *Bombus hyperboreus*) would differ from those of truly social bumblebees. We also hypothesised that the eye properties of species regularly occurring in forested environments would differ from those restricted to open landscapes. To characterise this, we generated two habitat metrics based on available literature: the *presence in forest* and the *forest affinity index*.

Importantly, the three variables *parasitism*, *presence in forest and forest affinity index* were largely related to phylogenetic relationships, with a typical heritability of 52%, 71% and 49% respectively. This link between phylogenetic and ecological variables does not necessarily undermine the efforts to study and disentangle ecological from phylogenetical drivers of visual traits (Morrissey and Ruxton, 2018), but does underline the absolute necessity to compute phylogenetically-controlled models in the following analyses.

#### The effects of social parasitism on visual traits

To test the hypothesis that the compound eyes of cuckoo bumblebees differ from those of truly social bumblebees, we quantified the effect of *parasitism* on IO angle, facet number, eye parameter and facet diameter. We focused on queens due to the low number of parasitic males in the dataset (N = 2) and found that they seem to differ from true queens in their eye size *(**Figure 2c, Figure S1**)*. In order to disentangle the effects of *parasitism* and eye size, we ran multiple regressions including both parameters. To compare models and effects, we use the Bayesian p-value (*p_MCMC_*) of each effect and the Deviance Information Criterion (*DIC*). We found significant effects of *parasitism* on IO angle, and eye parameter *(**Figure 4**, **Table 2**, **Table S6**)*. In particular, the eye parameter was best explained by models including the effects of *parasitism*, eye size and of their interaction, indicating that part of the negative effect of *parasitism* on the eye parameter was not because parasitic queens had slightly smaller eyes. Phylogenetic heritability did not change substantially by including the effects of *parasitism (**Table S5**, **Table S6**)*, suggesting that the effect of *parasitism* is not mediated by phylogeny. Overall, these findings indicate that parasitic queens have a lower eye parameter than truly social bumblebees. Whether this is also the case in parasitic males remains to be tested with more individuals, but the relatively low eye parameters in males of the parasitic species *Bombus barbutellus* and *Bombus rupestris* suggest that this might be the case *(**Figure 2c**)*.

**Figure 4:**
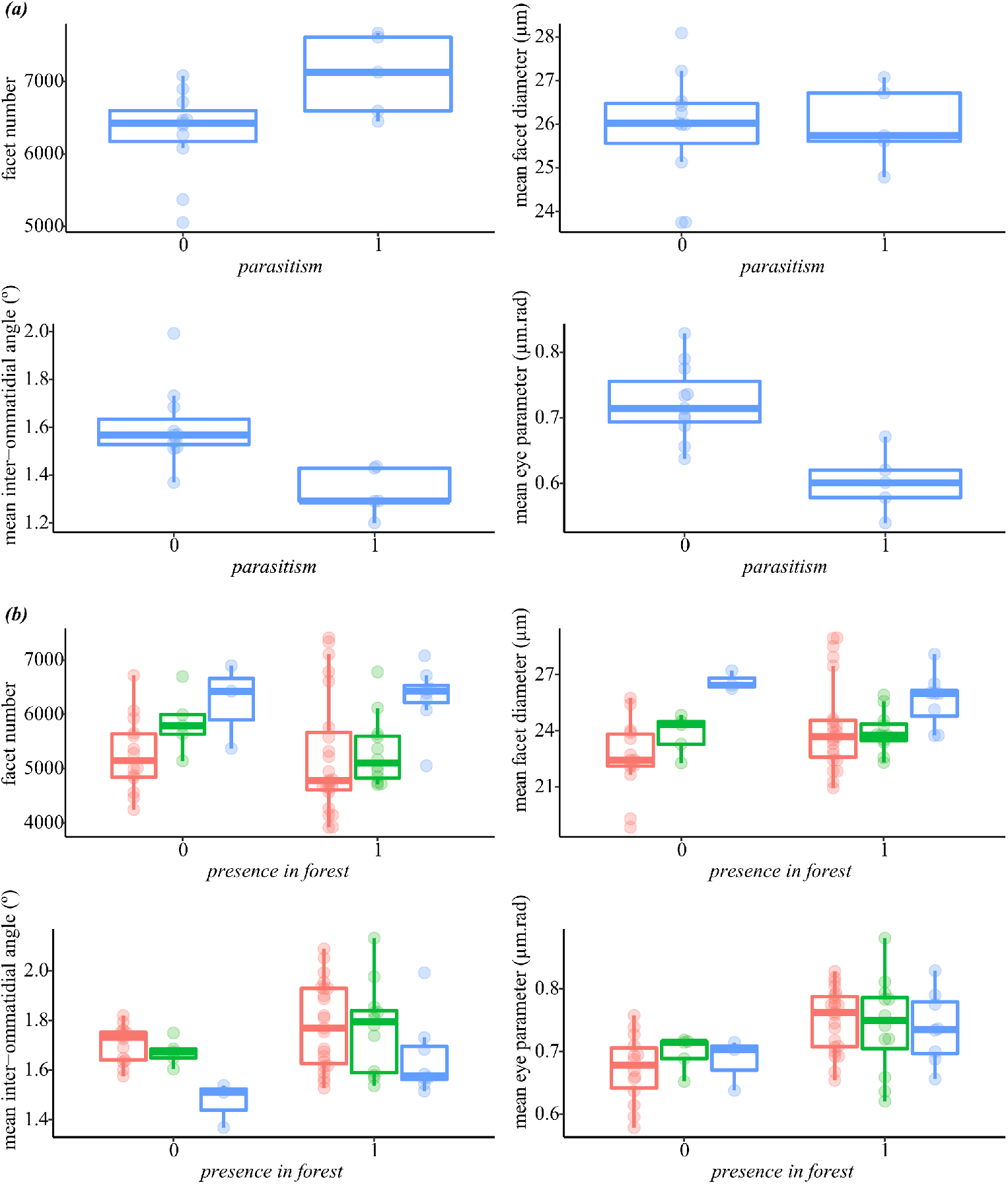
Effects of ecological variables on the main visual traits. *(a)* Effect of social *parasitism* on visual traits in bumblebee queens. *(a)* Effect of *presence in forest* on visual traits in bumblebee workers (red), males (green) and queens (blue).

**Table 2:** Summary of the effects of *parasitism* on visual traits in queens. Simple or multiple Bayesian linear regressions including *parasitism* and/or eye size were modelled on each visual trait. The lowest Deviance Information Criterion (*DIC*) indicates the best model for each trait (highlighted in beige). Significant positive or negative effects are reported in blue or red (respectively). The significance was based on the Bayesian p-value (*p_MCMC_*<0.05). The NULL model only included phylogeny and replication as random variables.

|  | Null | *parasitism* |  | eye size |  | *parasitism* and eye size |  | *parasitism* and eye size and their interaction |  |
| --- | --- | --- | --- | --- | --- | --- | --- | --- | --- |
|  | DIC | DIC | significant effects | DIC | significant effects | DIC | significant effects | DIC | significant effects |
| mean facet diameter ($\mu\text{m}$) | 37.11 | 37.54 | | 4.87 | eye size | 5.27 | eye size | -10.62 | eye size |
| mean IO angle ($^{\circ}$) | 15.69 | 20.66 | parasitism | 24.94 | eye size | 25.06 | parasitism<br>eye size | 18.01 | parasitism<br>eye size |
| facet number | -34.33 | -32.30 |  | 3.64 | eye size | 4.51 | eye size | -10.02 | eye size |
| mean eye parameter ($\mu\text{m}.\text{rad}$) | 25.37 | 30.04 | parasitism | 30.16 | | 32.04 | parasitism | 23.16 | parasitism |

To our knowledge, this is the first time that differences in the visual system of parasitic and non-parasitic bumblebees have been described. Essentially, cuckoo bumblebees favour optical resolution at the cost of light sensitivity. Are there evolutionary mechanisms for these differences? We identified several possible explanations:

##### (1) Reduced foraging demands

In social bumblebees, queens need to be highly efficient foragers when founding their colony (Goulson, 2010), leading to a high selective pressure on solitary founding (Harpur et al., 2017), that likely applies in part to sensory traits essential to foraging such as vision and olfaction. This constraint on foraging is likely to be weaker in parasitic queens that gather nectar only for their own consumption (with the exception of *B. hyperborreus* (Hines and Cameron, 2010)). This might explain why the visual traits of cuckoo bumblebees differ from true bumblebees, although it is unclear why a smaller eye parameter would be advantageous for the ecology of parasitic bumblebees.

##### (2) Simpler navigation tasks

Similarly, cuckoo queens likely perform few navigation tasks as they are not central place foragers. To navigate, bees rely on a light-sensitive Dorsal Rim Area (DRA) at the top of the compound eye. In cuckoo queens, weaker navigation requirements may have caused a reduction of the DRA and thus decreased the mean light sensitivity of the eye. However, this effect is unlikely to impact the mean eye parameter given that the DRA represents only a very small portion of the eye.

##### (3) Habitat-mediated effect

The effect of *parasitism* may simply be mediated by the habitat in which cuckoo bees live (seemingly more open than other bees, which is expected to have similar negative effects on the eye parameter). However, in a model including the effects of *presence in forest*, *parasitism* and eye size, the effect of parasitism was still negative although not significant, probably due to the large number of explanatory variables. Moreover, there was no obvious consistent differences of eye parameters between parasitic queens present in open (*B. hyperborreus, B. rupestris*) or forested habitats (*B. campestris, B. bohemicus*). This cause is thus unlikely.

##### (4) Visually controlled host search

A critical step in the lifecycle of parasitic queens is locating the hive of their host species. While numerous studies have demonstrated the importance of olfactory cues from the host species to achieve this (reviewed in Lhomme and Hines, 2018), it is unknown if parasitic queens use visual cues to locate the host nest. At a small range, it is possible that the nest entrance stands out as a small dark spot in the eyes of the parasite (as is the case in stingless bees (Biesmeijer et al., 2005; Tichit et al., 2020)). In this case, investing in improved resolution could be advantageous for locating hives. This is specifically the case if the search occurs during the brightest hours of the day and/or in open micro-habitats, which seems to be the case in *Bombus rupestris* (personal observation). Furthermore, high resolution could be beneficial in the hypothetical case where parasitic queens use visual cues from the host bumblebees themselves entering and leaving the hive (in addition to olfactory cues (Cederberg, 1983)) to guide them. While current lack of information makes it difficult to assess the validity of this explanation, it does highlight the need to investigate whether cuckoo bees use vision during host search.

##### (5) Lower flight speed

Naturalists are familiar with the low-frequency buzz of flying cuckoo bumblebees (Lhomme and Hines, 2018). This is because cuckoo bumblebees seemingly have slower flight than other bumblebees (Lhomme and Hines, 2018; Mossberg and Cederberg, 2012), although there are no documented measurements of flight speed in these species. Low flight speeds generate less motion blur than higher speeds and this may potentially enable cuckoo bumblebees to shift the trade-off of resolution and sensitivity towards greater resolution (as is the case in walking insects (Snyder, 1979)). This factor is thus likely to contribute to the low eye parameter of parasitic queens. More generally, variations of typical flight speeds across species of flying insects may generate a fine tuning of visual traits (as recently suggested (Baird et al., 2020)) such as the eye parameter.

##### (6) Pleiotropy and side-effects

Other factors may explain the unusual eye properties of parasitic queens without involving direct selection of visual traits. Cuckoo bees evolved a number of anatomical features such as a thicker cuticle, larger mandibles and associated muscles, likely as adaptations to their parasitic lifestyle (Lhomme and Hines, 2018). These features may indirectly impact the visual system via pleiotropic or morphological effects on the eye anatomy. For instance, the massive mandibles of parasitic queens may constrain the size and shape of their compound eyes, impacting their optics and thus probably their function. More generally, this underlines the need for visual ecologists to consider possible correlations between head and eye anatomy, e.g. between face length and eye shape in bees.

To conclude, there are several plausible explanations for the link between social parasitism and visual traits. We think that it is thus reasonable to move from correlation to causation by proposing that social parasitism has driven a shift of visual traits in bumblebees. To explore the evolutionary mechanisms discussed above, researchers would benefit from including more parasitic species in their comparative studies, and in particular more males, because on the contrary to parasitic queens, they have a life history similar to the males of other bumblebees.

#### The effects of forest habitats on visual traits

In comparison to open habitats, forests generate specific visual conditions (for instance lower light intensities, occlusion of part of the horizon and the sky) that could constraint the visual systems of bumblebees. To test the hypothesis that visual traits of species regularly occurring in forested environments would differ from those restricted to open landscapes, we investigated the effects of the *presence in forest* on eye properties in worker bumblebees (where there is no confounding effect of parasitism and the number of samples is highest (N = 37)). We followed a procedure similar to that used for *parasitism* by comparing the null models to simple and multiple regressions including *presence in forest* and/or eye size as explanatory variables. Facet number, facet diameter and eye parameter were best explained by models including the effects of *presence in forest*, eye size and of their interaction *(**Figure 5**, **Table 3a, Table S7**)*. In particular, there was a consistent positive effect of the *presence in forest* on the eye parameter, and this effect persisted even when the interaction of eye size and *presence in forest* was considered. This indicates that, even at constant eye size, workers of species occurring in forests had higher eye parameters than those that occur primarily in open landscapes.

**Table 3:** Summary of the effects of habitat *(presence in forest (a)* and *forest affinity index (b))* on visual traits in workers. Simple or multiple Bayesian linear regressions including habitat and/or eye size were modelled on each visual trait. The lowest Deviance Information Criterion (*DIC*) indicates the best model for each trait (highlighted in beige). Significant positive or negative effects are reported in blue or red (respectively). The significance was based on the Bayesian p-value (*p_MCMC_* <0.05). The NULL model only included phylogeny and replication as random variables.

| (a) | Null | *presence in forest* |  | eye size |  | *presence in forest* and eye size |  | *presence in forest* and eye size and their interaction |  |
| --- | --- | --- | --- | --- | --- | --- | --- | --- | --- |
|  | DIC | DIC | significant effects | DIC | significant effects | DIC | significant effects | DIC | significant effects |
| mean facet diameter ($\mu\text{m}$) | 91.44 | 91.40 | | 14.28 | eye size | 12.23 | forest<br>eye size | 3.93 | eye size<br>interaction |
| mean IO angle ($^\circ$) | 59.45 | 59.19 | | 41.98 | eye size | 41.85 | forest<br>eye size | 45.23 | forest<br>eye size |
| facet number | 82.83 | 83.70 |  | 16.59 | eye size | 14.43 | forest<br>eye size | 9.17 | forest<br>eye size |
| mean eye parameter ($\mu\text{m}.\text{rad}$) | 93.06 | 88.87 | forest | 69.29 | eye size | 69.27 | forest<br>eye size | 67.58 | forest<br>eye size |

| (b) | Null | *forest affinity index* |  | eye size |  | *forest affinity index* and eye size |  | *forest affinity index* and eye size and their interaction |  |
| --- | --- | --- | --- | --- | --- | --- | --- | --- | --- |
|  | DIC | DIC | significant effects | DIC | significant effects | DIC | significant effects | DIC | significant effects |
| mean facet diameter ($\mu\text{m}$) | 91.44 | 86.56 | forest | 14.28 | eye size | 14.67 | eye size | 14.00 | eye size |
| mean IO angle ($^\circ$) | 59.45 | 59.76 | | 41.98 | eye size | 42.93 | eye size | 39.27 | eye size |
| facet number | 82.83 | 83.72 |  | 16.59 | eye size | 16.45 | eye size | 16.34 | eye size |
| mean eye parameter ($\mu\text{m}.\text{rad}$) | 93.06 | 90.85 | forest | 69.29 | eye size | 70.50 | eye size | 64.59 | forest<br>eye size<br>interaction |

The effect of forest seems particularly pronounced in *Bombus transversalis* that lives in tropical rainforests (Olesen, 1989) (note that the effects of *presence in forest* on visual traits were the same when *B. transversalis* was excluded from the dataset ***Table S8**)*. The workers of this species reached “queen-sized” eyes, with a strong investment in increased facet diameter, facet number and eye parameter *(**Figure 2b-c**)*. The large eyes of *B. transversalis* were also correlated to an unusually large body (inter-tegular width = 6.0 ± 0.4 mm, N = 4). This appears to be consistent with the suggestion made for nocturnal bees that a large body size may facilitate a shift towards challenging visual environments (in this case a tropical rainforest) by providing larger eyes to a bee (Dorey et al., 2020).

To assess whether our findings were sensitive to the variables chosen to reflect the association of species with forests, we explored whether the *forest affinity index* (mostly based on a published review of bumblebee habitat (Liczner and Colla, 2019), and thus mostly independent from the *presence in forest*) had the same effects on the visual traits of workers. We found significant positive effects of *forest affinity index* on IO angle and on the eye parameter *(**Figure S2**, **Table 3b**, **Table S9**)*. The results on the eye parameter show a good agreement between the two forest variables, which reinforces our conclusion regarding the positive correlation between forest and this visual trait.

To further evaluate the consistency of the effect of the *presence in forest* and *forest affinity index* on visual traits, we performed a similar analysis in males and queens of truly social species (to avoid confounding effects of *parasitism*). In both castes, the two forest variables had no significant effect on the visual traits that were generally best explained by phylogeny only (null model) or by eye size *(**Table S5**, **Table S7**)*. This lack of clear effect of the forest variables in castes other than workers could be explained by the low explanatory power of the models due to the small number of specimens in males and queens. Alternatively, this could reflect biological differences. Despite sharing the same species-specific habitat, the three castes interact differently with it. These interactions are under different selective pressures that may interfere with the direct habitat effects. For instance, male fitness is tightly linked to mating success. Moreover, the larger compound eyes of males and queens *(**Figure 2**)* may perform well enough to cope with forests without further modifications. On the contrary, workers have small eyes that may benefit from the tuned eye properties described above.

It is important to note that it was necessary to simplify the scoring of forest variables, because the habitat preferences of bumblebee species have rarely been quantified. For example, with the first metric *presence in forest*, we reported any link of species with forests regardless of its strength or the type of forest habitat. There was no distinction between *B. balteatus* that occasionally visits sparse forest dominated by small arctic birches (Ranta and Lundberg, 1981), and *B. tranversalis* that lives in the dense tropical rainforest (Olesen, 1989). More quantitative information about the differences between the visual habitats occupied by different bumblebee species is necessary to better understand if and how these shapes visual traits.

Our results suggest that the association with forest habitats is linked to the visual properties of bumblebee compound eyes. The higher eye parameter of species associated with forests indicates a higher investment towards improved sensitivity, at the cost of resolution. We also find that the extent of the field of view (FOV) tends to be larger in species living in forests *(**Figure 2a-c**)*. Because the direction of the crystalline cones greatly determines the exact shape of the FOV (Stavenga, 1979), this possibility would need to be confirmed by investigating the inner anatomy of the compound eyes. We identified several selective pressures that the extent of forest in the habitats could impose on visual traits, and more specifically on the eye parameter:

##### (1) Higher motion blur

In forests, the proximity of obstacles increases the optical motion blur experienced by flying bumblebees, which in turn diminishes their ability to resolve visual details. This may be detrimental if the bees are more likely to collide with obstacles, or less likely to detect flowers. Bumblebees may compensate this problem behaviourally by reducing their flight speed, as they do when entering artificial environments of increasing proximity (Linander et al., 2016). However, a sustained flight speed reduction would likely decrease the foraging efficiency of bumblebees in view of their wide foraging range (Westphal et al., 2006). Instead, forested habitats may select for durable adjustments of visual traits that offset the additional blur generated by the proximity of obstacles. For animals, higher motion blur is a visual challenge similar to lower light intensity (Snyder et al., 1977), which supports the higher eye parameter in forested habitat.

##### (2) Lower light intensity

Forests are typically dimmer than open landscapes. As for motion blur, a lower light intensity can be transiently compensated behaviourally via a reduction of flight speed (as when *B. terrestris* land in dim light (Reber et al., 2016)), with the cost of a poorer foraging efficiency. However, only adjustments of the visual system could permanently compensate for dim light in forests. a consistent presence in forest would thus select for higher eye parameter in bumblebees, which trades off sensitivity at the cost of resolution.

##### (3) Hardly discernible skyline

The compound eyes of bees seem adapted to perform optimal visual sampling (with highest resolution) around the skyline (Baumgärtner, 1928; Taylor et al., 2019) where visual information is highest (Cronin et al., 2014). Dense forests such as the tropical rain forest lack a visible skyline so there may be no selective advantage to improve optical resolution in forest species because it would not provide much additional visual information (as underlined in (Feller et al., 2020)). This, in turn, may contribute to increasing the eye parameter in bumblebee species occurring in forests.

Considering these potential selective mechanisms, we propose that the visual habitat, and particularly an association with forests, is a driver in the evolution of visual traits in bumblebees. To our knowledge, this is the first time that habitat-related (and likely habitat-driven) eye modifications have been described in bees. So far, only the effects of mating behaviours (e.g. Streinzer and Spaethe, 2014a) and activity period (e.g. Somanathan et al., 2009) on visual traits have been demonstrated. That visual habitat affects visual traits in bees is not surprising given that recent work has shown a relationship between visual traits and micro-habitats in fruit flies, damselflies and aphids (Farnier et al., 2015; Ian W Keesey et al., 2020; Scales and Butler, 2016). Effects of forest on optical resolution of compound eyes were suggested in a meta-analysis across arthropods (Feller et al., 2020), while vertebrates living in forests seem to have higher light sensitivity than their counterparts living in open landscapes (Veilleux and Lewis, 2011). Our work adds to this body of knowledge and highlights the importance of considering the interplay between visual habitat and visual traits of species when trying to understand the factors driving their evolution, or to assess their ability to adapt to environmental changes.

## Conclusion

### Summary of the study

In the present study, we used high-throughput micro-CT techniques to explore the diversity of visual traits of compound eyes of bumblebees. Our results confirmed the prediction that visual traits vary substantially across bumblebee sizes, castes and species. We found that phylogenetic proximity poorly explained species-specific variations. Instead, these variations were better accounted for by two ecological factors: social parasitism and habitat. Social parasitic queens had lower eye parameters than other bumblebee queens, while workers of species associated with forested habitat had different visual traits, including higher eye parameters, than those living in open landscapes. We discuss why these modified visual traits towards improved resolution (respectively sensitivity) are likely to provide selective advantages to parasitic bumblebees (respectively bumblebees living in forests), and conclude that social parasitism and forest habitat may have been substantially driving the diversification of compound eyes in bumblebees, leading to the current variability of visual traits.

### Implications of the study for landscape and conservation ecology - “why visual traits matter”

Our study underlines the tight link between ecological variables and the structure and function of compound eyes in bumblebees. In particular, it highlights possible relationships between the habitat niche and the visual abilities of species. Given the importance of the visual system of bumblebees to perform key behaviours such as flying (Baird et al., 2020), foraging (Dafni et al., 2007), navigating (Mandal, 2018), and mating (Paxton, 2005), it is both valuable and necessary for functional ecologists to include visual properties in trait-based approaches. We identified several points where a consideration of visual traits would be fruitful:

#### (1) Visual traits as predictors of habitat preferences

If (as the present study suggests) the habitat niche drives the evolution of visual traits, the former can be used to predict the habitat requirements of animals whose distributions and ecologies are poorly known. In a recent study, Dorey et al. (Dorey et al., 2020) demonstrated that studying visual anatomy was sufficient for predicting the temporal niche, e.g. the nocturnality, of bees with unknown activity patterns. Similarly, the study of visual traits in understudied species may help to predict their habitat niche (for example if they are more associated with open landscapes or cluttered forests).

#### (2) Visual traits as determinants of spatial distributions

To investigate intrinsic drivers of the spatial distribution and ecological niche of bumblebees, ecologists generally consider physiological or life-history traits such as tongue length and nesting types (Persson et al., 2015). For example, tongue length affects the type of flowers that are profitably exploited by bumblebee species, and thus determines their foraging niche (Ranta and Lundberg, 1980), which may in turn influence bees’ spatial distribution. Recently, more attention has been given to habitat requirements and how they likely also play a major role to determine the spatial distribution of bumblebees (Liczner and Colla, 2020). Visual abilities of bumblebees may constraint both habitat (as the present study suggests) and foraging niche (given that vision is crucial to detect and discriminate flowers). It is thus important to examine the role of visual traits (and not only of non-sensory traits) in determining the spatial distribution of species.

#### (3) Visual traits in risk assessments of anthropogenic changes

Anthropogenic activities, in particular landscape changes, strongly impact bumblebees and their habitat (Le Féon et al., 2010). Under these changing conditions, bumblebees may experience a mismatch between their visual abilities and the transformed habitat. For example, we found that bumblebees associated with forest possess compound eyes tuned to obtain better sensitivity at the cost of resolution. This characteristic may become disadvantageous in a more open landscape created by human activities. In turn, this may affect visually guided behaviours in bumblebees with negative consequences on their fitness that would contribute to their decline. Climate change may also accentuate the mismatch (as described for other traits (Miller-Struttmann et al., 2015)) between bumblebee eyes and their environment, by affecting the environment and/or the eyes themselves. Increasing temperatures due to climate change could for instance increase the variability of body size in bumblebees (Kelemen and Dornhaus, 2018), with possible impacts on compound eye size and thus visual traits (due to the allometric relationship highlighted in numerous studies including the present work). Bumblebees may compensate the mismatch between their eyes and the transformed environment via plastic adjustments of their visually-guided behaviours, or even (as may be the case in other insects (Dudaniec et al., 2018)) via local climate-driven visual adaptations. Alternatively, vision-mediated impacts of anthropogenic changes may lead to population declines and even contribute to the extinction of bumblebee species. In order to identify sensitive species and better preserve these key pollinators, we encourage conservation ecologists to take into account visual traits of bumblebees in trait-based risk assessments of landscape change (Persson et al., 2015) and climate change (Sirois-Delisle and Kerr, 2018; Williams et al., 2009).

## Supporting information

Electronic Supplementary Material

## Acknowledgments

We would like to thank Björn Cederberg, Bo Svensson, Tobio Aarts, Jonno Stedler, Gavin Taylor, Göran Holmström, Cristiane Krug, Isabel Alves dos Santos, Maria Cristina Arias, Karin Johnson as well as the staff at the Abisko Scientific Research Station, the National Research Institute of Amazonas, the University of São Paulo and the biological museum at Lund University for help and advice during sample collection. We are grateful to Inga Tuminaite, Gavin Taylor, David Wilby, Marie Guiraud, Rajmund Mokso, Kazimir Wanelik and Viktor Håkansson for their support during micro-CT scanning at the Diamond Light Source, UK. Thanks to Viktor Håkansson, Julia Meneghello, Gavin Taylor, Emelie Karlsson and Alexander Woin for help during volumetric and computational analysis. We would like to thank Heather Hines for sharing data on bumblebee phylogeny from her original work. We are grateful to Charlie Cornwallis, Jessica Abbott and Philip Downing for advice regarding data exploration and statistical analysis.

## Author contributions

P.T and E.B. designed the study and carried out collection of samples. P.T. performed sample preparation. P.T., A.B., C.R. and E.B. carried out sample imaging. P.T. performed volumetric, computational and statistical analysis. P.T and E.B. drafted the manuscript. P.T., A.B., C.R. and E.B. critically revised the manuscript.

## Data availability

The original image data of all bee compound eyes analysed in this study will be made available on MorphoSource prior to publication. The raw data used to generate the topologies of bee eye properties will be available on Dryad. A report of the R analyses performed in this study will be provided as a supplementary material.

