## Supplementary material for "‘Same same but different’ Exploring evolutionary and ecological causes of eye diversification in bumblebees": Electronic Supplementary Material

### Supplementary Materials

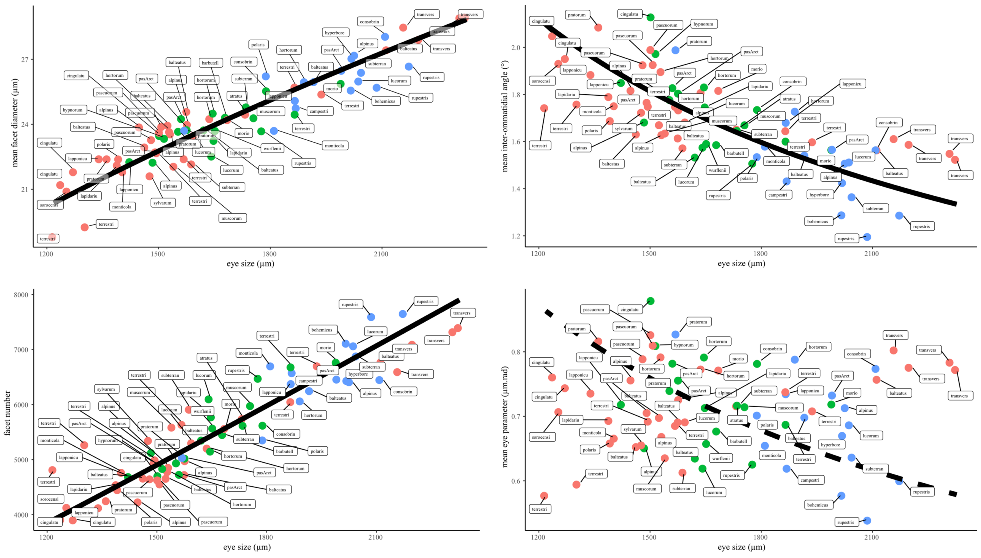


**Figure S1: Allometric relationship between eye size and the main visual traits**. Each point corresponds to an individual bumblebee: workers (red), males (green) and queens (blue). The plain (respectively dashed) black lines represent the significant (respectively non-significant) allometric relationship obtained by fitting a standardised major axis to the log-transformed eye size and visual trait.

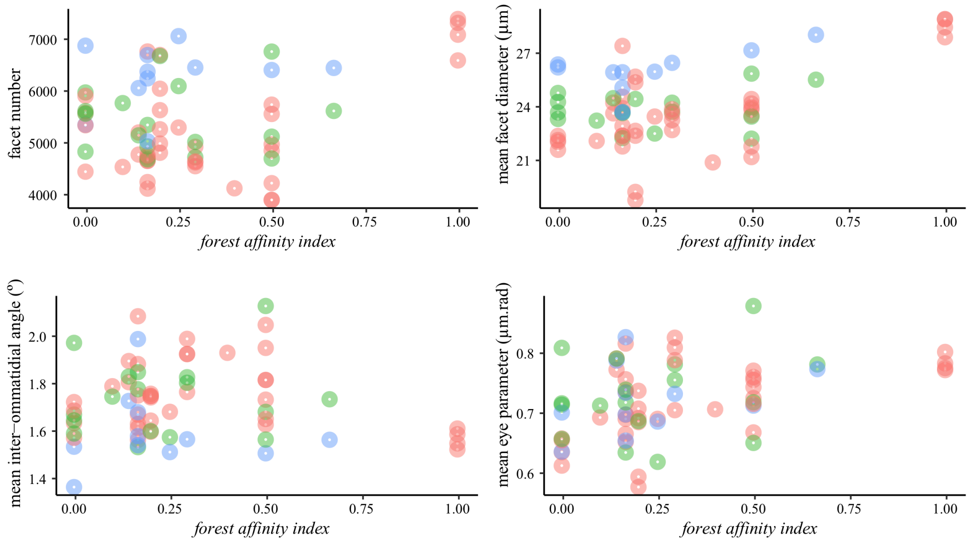


**Figure S2: Effects of the *forest affinity index* on the main visual traits in bumblebee workers (red), males (green) and queens (blue).**

**Table S1: Value of the visual traits for each bumblebee in the study (N = 72).** Note that the species label for *Bombus pascuorum smithianus* is 'pasArct', while 'pascuorum' refers to the subspecies *Bombus pascuorum pallidofacies*.

| mean facet diameter (µm) | mean facet area (µm²) | mean curvature (µm) | mean IO angle (°) | mean eye area (µm²) | caste | facet number | mean eye parameter (µm.rad) | extent of the FOV (%) | lowest eye parameter (µm.rad) | highest eye parameter (µm.rad) | species | highest facet diameter (µm) | lowest IO angle (°) | eye size |
|---|---|---|---|---|---|---|---|---|---|---|---|---|---|---|
| 23.58 | 483.73 | 884.09 | 1.63 | 2355546.61 | F | 4869.56 | 0.67 | 20.24 | 0.45 | 0.91 | alpinus | 25.80 | 1.18 | 1534.78 |
| 24.02 | 500.22 | 851.32 | 1.74 | 2496932.88 | F | 4991.69 | 0.73 | 23.13 | 0.50 | 0.99 | alpinus | 25.56 | 1.22 | 1580.17 |
| 23.91 | 496.58 | 806.28 | 1.82 | 2106998.31 | F | 4243.04 | 0.76 | 21.15 | 0.52 | 0.99 | alpinus | 25.88 | 1.31 | 1451.55 |

| | | | | | | | | | | | | | | |
|---|---|---|---|---|---|---|---|---|---|---|---|---|---|---|
| 22.28 | 430.74 | 814.32 | 1.69 | 2214213.71 | M | 5140.49 | 0.65 | 24.28 | 0.45 | 0.91 | alpinus | 24.15 | 1.20 | 1488.02 |
| 27.22 | 640.88 | 1120.64 | 1.51 | 4118166.41 | Q | 6425.84 | 0.71 | 23.66 | 0.47 | 1.00 | alpinus | 29.36 | 1.03 | 2029.33 |
| 27.47 | 659.59 | 1052.27 | 1.57 | 4474327.78 | F | 6783.49 | 0.76 | 24.38 | 0.48 | 1.12 | balteatus | 30.77 | 1.13 | 2115.26 |
| 23.98 | 499.92 | 891.77 | 1.67 | 2335328.77 | F | 4671.36 | 0.70 | 19.97 | 0.46 | 0.97 | balteatus | 25.71 | 1.12 | 1528.18 |
| 22.98 | 459.73 | 882.16 | 1.63 | 2149072.61 | F | 4674.64 | 0.65 | 19.32 | 0.41 | 0.89 | balteatus | 24.91 | 1.11 | 1465.97 |
| 24.62 | 525.93 | 928.13 | 1.62 | 2492637.68 | F | 4739.45 | 0.69 | 19.84 | 0.49 | 0.92 | balteatus | 26.51 | 1.18 | 1578.81 |
| 23.77 | 492.14 | 918.82 | 1.54 | 2640831.52 | M | 5366.01 | 0.64 | 20.40 | 0.47 | 0.80 | balteatus | 26.12 | 1.21 | 1625.06 |
| 25.99 | 588.73 | 1029.92 | 1.54 | 3687845.56 | Q | 6264.09 | 0.70 | 23.44 | 0.46 | 0.90 | balteatus | 28.89 | 1.10 | 1920.38 |
| 24.35 | 517.83 | 914.19 | 1.59 | 2831283.30 | M | 5467.58 | 0.68 | 23.89 | 0.46 | 0.91 | barbutell | 27.34 | 1.19 | 1682.64 |
| 25.61 | 570.08 | 1161.55 | 1.29 | 4081887.19 | Q | 7126.47 | 0.58 | 19.17 | 0.43 | 0.76 | bohemicus | 28.12 | 1.06 | 2020.37 |
| 24.79 | 532.08 | 1048.01 | 1.44 | 3506926.80 | Q | 6590.99 | 0.62 | 20.98 | 0.44 | 0.84 | campestri | 26.73 | 1.08 | 1872.68 |
| 21.24 | 392.12 | 620.94 | 2.05 | 1537814.60 | F | 3921.83 | 0.76 | 23.96 | 0.53 | 1.05 | cingulatu | 23.04 | 1.53 | 1240.09 |
| 21.84 | 414.74 | 694.92 | 1.96 | 1624142.71 | F | 3916.08 | 0.75 | 22.25 | 0.45 | 1.03 | cingulatu | 24.21 | 1.29 | 1274.42 |
| 23.51 | 480.50 | 687.50 | 2.13 | 2267082.68 | M | 4718.22 | 0.88 | 28.02 | 0.51 | 1.35 | cingulatu | 26.11 | 1.37 | 1505.68 |
| 25.57 | 570.26 | 891.13 | 1.74 | 3214288.62 | M | 5636.50 | 0.78 | 23.91 | 0.49 | 1.16 | consobrin | 28.47 | 1.23 | 1792.84 |
| 28.09 | 690.12 | 1072.41 | 1.57 | 4462386.57 | Q | 6466.07 | 0.78 | 23.04 | 0.50 | 1.10 | consobrin | 31.50 | 1.15 | 2112.44 |
| 23.69 | 489.89 | 753.80 | 1.90 | 2348653.15 | F | 4794.24 | 0.79 | 25.30 | 0.49 | 1.14 | hortorum | 26.52 | 1.32 | 1532.53 |
| 24.29 | 515.46 | 812.93 | 1.81 | 2691104.02 | F | 5220.80 | 0.77 | 25.70 | 0.48 | 1.15 | hortorum | 27.06 | 1.29 | 1640.46 |
| 25.99 | 590.57 | 904.71 | 1.73 | 3589777.38 | Q | 6078.45 | 0.79 | 26.47 | 0.49 | 1.07 | hortorum | 29.33 | 1.23 | 1894.67 |
| 24.54 | 526.80 | 799.39 | 1.83 | 2721336.68 | M | 5165.80 | 0.79 | 26.72 | 0.51 | 1.15 | hortorum | 27.53 | 1.35 | 1649.65 |
| 27.07 | 634.04 | 1162.06 | 1.43 | 4089125.72 | Q | 6449.28 | 0.67 | 21.00 | 0.45 | 0.86 | hyperbore | 29.74 | 1.01 | 2022.16 |
| 23.38 | 475.66 | 731.40 | 1.98 | 2307087.69 | M | 4850.27 | 0.81 | 27.57 | 0.47 | 1.21 | hypnorum | 25.95 | 1.29 | 1518.91 |
| 22.44 | 436.40 | 718.17 | 1.89 | 1805178.25 | F | 4136.50 | 0.74 | 21.10 | 0.53 | 0.98 | lapponicu | 24.03 | 1.40 | 1343.57 |
| 22.30 | 432.11 | 730.96 | 1.85 | 2032341.03 | M | 4703.26 | 0.72 | 26.34 | 0.50 | 0.92 | lapponicu | 24.54 | 1.32 | 1425.60 |
| 25.13 | 547.10 | 944.14 | 1.68 | 3497495.43 | Q | 6392.77 | 0.74 | 26.70 | 0.48 | 1.09 | lapponicu | 27.00 | 1.11 | 1870.16 |
| 22.15 | 425.30 | 742.67 | 1.79 | 1937441.90 | F | 4555.49 | 0.69 | 21.28 | 0.47 | 0.94 | lapidariu | 23.94 | 1.31 | 1391.92 |
| 23.28 | 471.43 | 798.82 | 1.75 | 2728330.37 | M | 5787.40 | 0.71 | 28.95 | 0.48 | 1.00 | lapidariu | 26.24 | 1.28 | 1651.77 |
| 23.51 | 480.33 | 840.51 | 1.69 | 2553695.72 | F | 5316.52 | 0.69 | 22.61 | 0.46 | 0.93 | lucorum | 25.54 | 1.20 | 1598.03 |
| 22.56 | 442.88 | 873.81 | 1.58 | 2708382.06 | M | 6115.43 | 0.62 | 22.29 | 0.43 | 0.86 | lucorum | 24.59 | 1.12 | 1645.72 |
| 26.02 | 588.18 | 1043.21 | 1.52 | 4163938.92 | Q | 7079.42 | 0.69 | 25.20 | 0.46 | 0.93 | lucorum | 28.37 | 1.07 | 2040.57 |
| 21.84 | 413.77 | 763.23 | 1.75 | 1977934.85 | F | 4780.26 | 0.67 | 22.50 | 0.46 | 0.88 | monticola | 23.42 | 1.25 | 1406.39 |
| 23.74 | 490.06 | 921.68 | 1.58 | 3290616.70 | Q | 6714.66 | 0.66 | 26.46 | 0.41 | 0.90 | monticola | 25.98 | 1.09 | 1814.01 |
| 24.24 | 511.00 | 790.11 | 1.82 | 2850318.66 | F | 5577.94 | 0.77 | 25.54 | 0.52 | 1.00 | morio | 26.82 | 1.37 | 1688.29 |
| 25.91 | 585.83 | 1027.23 | 1.57 | 3972794.94 | M | 6781.51 | 0.72 | 25.15 | 0.38 | 1.11 | morio | 29.36 | 0.93 | 1993.19 |
| 22.09 | 425.58 | 785.87 | 1.64 | 2383975.55 | F | 5601.69 | 0.64 | 20.32 | 0.43 | 0.81 | muscorum | 24.53 | 1.29 | 1544.01 |
| 24.33 | 516.27 | 844.97 | 1.67 | 3093939.35 | M | 5992.82 | 0.72 | 26.88 | 0.50 | 0.92 | muscorum | 27.61 | 1.35 | 1758.96 |
| 22.75 | 452.05 | 763.65 | 1.77 | 2234905.81 | F | 4943.95 | 0.71 | 22.46 | 0.46 | 0.92 | pasArct | 25.26 | 1.31 | 1494.96 |
| 23.78 | 495.27 | 786.31 | 1.81 | 2495434.62 | M | 5038.52 | 0.76 | 27.26 | 0.46 | 1.01 | pasArct | 27.20 | 1.28 | 1579.69 |
| 24.29 | 516.00 | 797.41 | 1.83 | 2447568.06 | M | 4743.34 | 0.78 | 25.97 | 0.47 | 1.10 | pasArct | 27.81 | 1.29 | 1564.47 |
| 26.52 | 614.71 | 1006.49 | 1.57 | 3979656.03 | Q | 6474.03 | 0.73 | 22.89 | 0.48 | 1.02 | pasArct | 29.58 | 1.13 | 1994.91 |
| 23.32 | 472.44 | 720.17 | 1.93 | 2202311.81 | F | 4661.59 | 0.79 | 23.81 | 0.51 | 1.02 | pascuorum | 26.05 | 1.42 | 1484.02 |
| 23.66 | 487.93 | 707.26 | 1.99 | 2265241.11 | F | 4642.53 | 0.83 | 25.84 | 0.54 | 1.08 | pascuorum | 26.57 | 1.50 | 1505.07 |
| 23.93 | 499.87 | 745.35 | 1.93 | 2284344.83 | F | 4569.92 | 0.81 | 23.58 | 0.50 | 1.07 | pascuorum | 26.81 | 1.36 | 1511.40 |
| 24.49 | 523.67 | 864.27 | 1.66 | 3015202.15 | F | 5757.80 | 0.72 | 20.84 | 0.46 | 0.96 | atratus | 27.06 | 1.27 | 1736.43 |
| 22.43 | 436.32 | 817.33 | 1.69 | 1947824.84 | F | 4464.21 | 0.66 | 20.56 | 0.45 | 0.90 | polaris | 24.10 | 1.16 | 1395.64 |
| 26.26 | 598.27 | 1038.73 | 1.54 | 3212105.00 | Q | 5369.01 | 0.70 | 19.74 | 0.50 | 0.93 | polaris | 28.33 | 1.14 | 1792.23 |
| 22.43 | 436.75 | 649.89 | 2.09 | 1862307.93 | F | 4264.01 | 0.82 | 25.51 | 0.53 | 1.07 | pratorum | 25.01 | 1.49 | 1364.66 |

| | | | | | | | | | | | | | | |
|---|---|---|---|---|---|---|---|---|---|---|---|---|---|---|
| 23.71 | 489.32 | 802.23 | 1.78 | 2422852.23 | M | 4951.44 | 0.74 | 24.25 | 0.49 | 1.07 | pratorum | 26.42 | 1.28 | 1556.55 |
| 23.75 | 489.80 | 709.75 | 1.99 | 2473960.76 | Q | 5050.95 | 0.83 | 25.62 | 0.57 | 1.09 | pratorum | 26.00 | 1.50 | 1572.88 |
| 25.74 | 573.98 | 1278.23 | 1.20 | 4367803.70 | Q | 7609.71 | 0.54 | 19.10 | 0.40 | 0.71 | rupestris | 28.16 | 0.93 | 2089.93 |
| 23.72 | 488.23 | 937.70 | 1.51 | 3168220.47 | M | 6489.15 | 0.63 | 25.65 | 0.46 | 0.83 | rupestris | 25.51 | 1.18 | 1779.95 |
| 26.71 | 617.75 | 1249.36 | 1.29 | 4736306.73 | Q | 7667.01 | 0.60 | 21.63 | 0.45 | 0.80 | rupestris | 29.09 | 0.98 | 2176.31 |
| 20.94 | 381.28 | 645.15 | 1.93 | 1579786.35 | F | 4143.39 | 0.71 | 22.84 | 0.50 | 0.94 | soroeensi | 22.81 | 1.47 | 1256.90 |
| 22.20 | 427.45 | 829.92 | 1.58 | 2533569.75 | F | 5927.14 | 0.61 | 19.85 | 0.43 | 0.82 | subterran | 24.59 | 1.24 | 1591.72 |
| 24.83 | 536.55 | 895.68 | 1.65 | 3022783.69 | M | 5633.72 | 0.72 | 24.41 | 0.50 | 0.97 | subterran | 27.19 | 1.26 | 1738.62 |
| 26.43 | 608.22 | 1142.13 | 1.37 | 4193811.29 | Q | 6895.23 | 0.64 | 18.51 | 0.43 | 0.87 | subterran | 29.38 | 1.04 | 2047.88 |
| 21.65 | 408.64 | 742.15 | 1.73 | 2190375.18 | F | 5360.13 | 0.66 | 22.39 | 0.44 | 0.87 | sylvarum | 24.11 | 1.28 | 1479.99 |
| 25.75 | 576.03 | 956.07 | 1.65 | 3493440.71 | F | 6064.66 | 0.74 | 26.28 | 0.49 | 1.00 | terrestri | 28.06 | 1.17 | 1869.07 |
| 25.42 | 561.10 | 967.37 | 1.60 | 3768524.30 | F | 6716.30 | 0.71 | 26.64 | 0.49 | 0.92 | terrestri | 27.63 | 1.13 | 1941.27 |
| 22.71 | 448.17 | 795.46 | 1.75 | 2244223.94 | F | 5007.50 | 0.69 | 23.94 | 0.45 | 0.93 | terrestri | 24.63 | 1.20 | 1498.07 |
| 22.44 | 437.05 | 782.08 | 1.75 | 2469850.15 | F | 5651.13 | 0.69 | 26.14 | 0.46 | 0.94 | terrestri | 24.15 | 1.23 | 1571.58 |
| 18.84 | 307.56 | 683.47 | 1.75 | 1485351.81 | F | 4829.50 | 0.58 | 22.92 | 0.32 | 0.89 | terrestri | 20.84 | 1.05 | 1218.75 |
| 19.30 | 322.76 | 683.15 | 1.76 | 1705036.99 | F | 5282.71 | 0.60 | 24.17 | 0.35 | 0.87 | terrestri | 21.33 | 1.12 | 1305.77 |
| 24.49 | 522.00 | 964.16 | 1.60 | 3495626.58 | M | 6696.65 | 0.69 | 28.29 | 0.40 | 1.00 | terrestri | 26.92 | 1.00 | 1869.66 |
| 28.95 | 728.71 | 1128.70 | 1.55 | 5345493.26 | F | 7335.60 | 0.78 | 27.34 | 0.53 | 1.09 | transvers | 31.90 | 1.15 | 2312.03 |
| 28.97 | 730.72 | 1140.29 | 1.53 | 5414829.77 | F | 7410.25 | 0.77 | 27.83 | 0.52 | 1.04 | transvers | 32.17 | 1.13 | 2326.98 |
| 27.95 | 682.01 | 1051.41 | 1.59 | 4847282.04 | F | 7107.33 | 0.78 | 28.38 | 0.53 | 1.04 | transvers | 30.80 | 1.20 | 2201.65 |
| 28.51 | 706.55 | 1053.43 | 1.62 | 4670701.68 | F | 6610.55 | 0.80 | 27.34 | 0.56 | 1.08 | transvers | 31.30 | 1.24 | 2161.18 |
| 23.74 | 490.87 | 901.28 | 1.59 | 2740964.00 | M | 5583.89 | 0.66 | 22.04 | 0.45 | 0.92 | wurflenii | 26.29 | 1.19 | 1655.59 |

**Table S2: Value of the visual traits for each other bees used for comparison in the study.**

| species | mean Facet diameter ($\mu m$) | mean Facet area ($\mu m^2$) | mean Curvature ($\mu m$) | mean IO angle (°) | mean eye area ($\mu m^2$) | caste | facet number | mean Eye parameter ($\mu$m.rad) | extent of the FOV (%) | lowest eye parameter ($\mu$m.rad) | highest eye parameter ($\mu$m.rad) | highest facet diameter ($\mu m$) | lowest IO angle (°) |
|---|---|---|---|---|---|---|---|---|---|---|---|---|---|
| *Apis mellifera* | 22.39 | 433.86 | 815.09 | 1.69 | 2440463.26 | F | 5624.99 | 0.66 | 27.59 | 0.42 | 0.93 | 1.15 | 24.62 |
| *Apis mellifera* | 22.52 | 439.06 | 849.23 | 1.67 | 2604449.57 | F | 5931.94 | 0.66 | 27.97 | 0.41 | 0.97 | 1.08 | 24.43 |
| *Scaptotrigona depilis* | 17.94 | 278.57 | 428.77 | 2.55 | 1159934.13 | F | 4163.95 | 0.80 | 38.32 | 0.57 | 1.08 | 1.82 | 19.33 |
| *Scaptotrigona depilis* | 18.10 | 283.34 | 446.39 | 2.47 | 1194947.05 | F | 4217.43 | 0.78 | 37.40 | 0.53 | 1.06 | 1.72 | 19.58 |
| *Scaptotrigona depilis* | 18.38 | 292.30 | 437.76 | 2.56 | 1142771.74 | F | 3909.54 | 0.82 | 35.81 | 0.59 | 1.11 | 1.84 | 19.57 |
| *Xylocopa tenuiscapa* | 31.55 | 863.57 | 1461.78 | 1.29 | 9960129.32 | Q | 11533.65 | 0.71 | 31.04 | 0.53 | 0.94 | 1.02 | 33.82 |
| *Megalopta genalis* | | | | | | Q | | | | 0.90 | 1.20 | 1.40 | 36.00 |

**Tables S3: Summary of the visual traits of the psyllids (a) and damselflies (b) used for comparison in the study.** Values were obtained from two previous studies (Farnier et al., 2015; Scales and Butler, 2016).

| *(a)* | N | mean | Standard deviation | minimum | 10% percentile | median | 90% percentile | maximum | Coefficient of variation (*CV*) | Relative half range (*HR*) |
|---|---|---|---|---|---|---|---|---|---|---|
| **facet number** | 4 | 217.50 | 70.42 | 160.00 | 169.00 | 195.00 | 284.00 | 320.00 | 32.37 | 36.78 |
| **highest facet diameter ($\mu$m)** | 4 | 12.38 | 1.25 | 11.00 | 11.30 | 12.25 | 13.55 | 14.00 | 10.10 | 12.12 |
| **eye size ($\mu$m)** | 4 | 0.17 | 0.05 | 0.14 | 0.14 | 0.15 | 0.21 | 0.24 | 27.93 | 28.25 |
| **eye parameter ($\mu$m.rad)** | 4 | 6.10 | 0.58 | 5.50 | 5.56 | 6.15 | 6.60 | 6.60 | 9.56 | 9.02 |
| **IO angle (º)** | 4 | 1.32 | 0.15 | 1.10 | 1.19 | 1.40 | 1.40 | 1.40 | 11.32 | 11.32 |

| *(b)* | N | mean | Standard deviation | minimum | 10% percentile | median | 90% percentile | maximum | Coefficient of variation (*CV*) | Relative half range (*HR*) |
|---|---|---|---|---|---|---|---|---|---|---|
| **lowest IO angle (º)** | 13 | 0.75 | 0.09 | 0.57 | 0.63 | 0.76 | 0.83 | 0.93 | 12.54 | 23.81 |
| **highest facet diameter ($\mu$m)** | 13 | 32.66 | 2.29 | 28.60 | 29.32 | 32.70 | 34.92 | 36.70 | 7.01 | 12.40 |

**Table S4: detailed results of the allometric study of visual traits.** Allometric curves and estimates were obtained by using the *SMATR* package (Warton et al., 2012). *b* and *α* are the estimated initial growth rate and scaling exponent of the allometric relationship respectively. Low and high CI correspond to the lower and upper limit of the confidence interval of the estimates.

| | ***b*** | ***b* low CI** | ***b* high CI** | ***α*** | ***α* low CI** | ***α* high CI** | **$R^2$** | **p value** |
|---|---|---|---|---|---|---|---|---|
| **mean facet diameter ($\mu$m)** | -0.35 | -0.51 | -0.19 | 0.54 | 0.49 | 0.59 | 0.85 | 0.00 |
| **mean IO angle (º)** | 2.48 | 2.11 | 2.85 | -0.70 | -0.82 | -0.59 | 0.53 | 0.00 |
| **facet number** | 0.24 | -0.08 | 0.56 | 1.09 | 0.99 | 1.19 | 0.85 | 0.00 |
| **mean curvature ($\mu$m)** | -0.52 | -0.86 | -0.19 | 1.07 | 0.98 | 1.18 | 0.84 | 0.00 |
| **mean eye parameter ($\mu$m.rad)** | 1.85 | 1.37 | 2.32 | -0.62 | -0.78 | -0.49 | 0.00 | 0.89 |
| **highest facet diameter ($\mu$m)** | -0.38 | -0.57 | -0.20 | 0.56 | 0.51 | 0.62 | 0.82 | 0.00 |
| **lowest IO angle (º)** | 2.32 | 1.90 | 2.75 | -0.70 | -0.84 | -0.58 | 0.36 | 0.00 |
| **lowest eye parameter ($\mu$m.rad)** | -2.33 | -2.80 | -1.85 | 0.62 | 0.49 | 0.79 | 0.01 | 0.40 |
| **highest eye parameter ($\mu$m.rad)** | 2.35 | 1.79 | 2.92 | -0.73 | -0.93 | -0.58 | 0.00 | 0.86 |
| **Extent of the FOV (%)** | -0.97 | -1.52 | -0.42 | 0.73 | 0.58 | 0.92 | 0.02 | 0.1 |

**Table S5: detailed results of the NULL models of the phylogenetic effects on visual traits in each caste.** The NULL model only included an intercept and phylogeny and replication as random variables. The phylogenetic heritability was the proportion of the posterior covariance explained by phylogeny. Low and high CI correspond to the lower and upper limit of the credible interval of phylogenetic heritability.

| WORKERS | DIC | phylogenetic heritability (%) | phylogenetic heritability low CI (%) | phylogenetic heritability high CI (%) |
|---|---|---|---|---|
| mean Facet diameter (µm) | 91.44 | 0.33 | 0.02 | 66.43 |
| mean IO angle (º) | 59.45 | 0.33 | 0.04 | 90.36 |
| facet number | 82.83 | 0.35 | 0.03 | 77.29 |
| mean Eye parameter (µm.rad) | 93.06 | 0.38 | 0.01 | 59.13 |

| MALES | DIC | phylogenetic heritability (%) | phylogenetic heritability low CI (%) | phylogenetic heritability high CI (%) |
|---|---|---|---|---|
| mean Facet diameter (µm) | 33.56 | 0.47 | 0.02 | 91.19 |
| mean IO angle (º) | -14.61 | 0.54 | 0.02 | 98.49 |
| facet number | 23.19 | 0.48 | 0.02 | 96.03 |
| mean Eye parameter (µm.rad) | 20.17 | 0.39 | 0.01 | 96.01 |

| QUEENS | DIC | phylogenetic heritability (%) | phylogenetic heritability low CI (%) | phylogenetic heritability high CI (%) |
|---|---|---|---|---|
| mean Facet diameter (µm) | 37.11 | 0.49 | 0.01 | 87.17 |
| mean IO angle (º) | 15.69 | 95.13 | 0.02 | 96.87 |
| facet number | -34.33 | 98.98 | 0.04 | 99.46 |
| mean Eye parameter (µm.rad) | 25.37 | 89.54 | 0.01 | 93.65 |

**Table S6: detailed results of the effects of parasitism on visual traits in queens.** Simple or multiple Bayesian linear regressions including parasitism and/or eye size were modelled on each visual trait. The lowest Deviance Information Criterion (*DIC*) indicates the best model for each trait (highlighted in beige). Significant positive or negative effects are reported in blue or red (respectively). The significance was based on the Bayesian p-value ($p_{MCMC}$ < 0.05). The phylogenetic heritability was the proportion of the posterior covariance explained by phylogeny.

Effects of *parasitism*

| | DIC | Slope | $p_{MCMC}$ | Phylogenetic heritability (%) |
|---|---|---|---|---|
| **mean facet diameter (µm)** | 37.54 | 0.11 | 0.87 | 0.46 |
| **mean IO angle (º)** | 20.66 | -1.13 | 0.05 | 0.49 |
| **facet number** | -32.30 | 0.81 | 0.18 | 0.55 |
| **mean eye parameter (µm.rad)** | 30.04 | -1.34 | 0.02 | 0.56 |

Effects of eye size

| | DIC | Slope | $p_{MCMC}$ | Phylogenetic heritability (%) |
|---|---|---|---|---|
| **mean facet diameter (µm)** | 4.87 | 0.87 | **0.00** | 0.67 |
| **mean IO angle (º)** | 24.94 | -0.65 | **0.00** | 0.53 |
| **facet number** | 3.64 | 0.72 | **0.00** | 0.54 |
| **mean eye parameter (µm.rad)** | 30.16 | -0.34 | 0.19 | 0.50 |

Effects of *parasitism* and eye size

| | DIC | ***parasitism* slope** | $p_{MCMC}$ | eye size slope | $p_{MCMC}$ |
|---|---|---|---|---|---|
| **mean facet diameter (µm)** | 5.27 | -0.41 | 0.38 | 0.91 | **0.00** |
| **mean IO angle (º)** | 25.06 | -0.87 | **0.03** | -0.58 | **0.00** |
| **facet number** | 4.51 | 0.40 | 0.31 | 0.67 | **0.00** |
| **mean eye parameter (µm.rad)** | 32.04 | -1.19 | **0.03** | -0.25 | 0.25 |

Effects of *parasitism* and eye size and of their interaction

| | DIC | ***parasitism* slope** | $p_{MCMC}$ | eye size slope | $p_{MCMC}$ | Interaction ***parasitism*** and eye size slope | $p_{MCMC}$ |
|---|---|---|---|---|---|---|---|
| **mean facet diameter (µm)** | -10.62 | -0.46 | 0.36 | 0.85 | **0.00** | 0.20 | 0.63 |
| **mean IO angle (º)** | 18.01 | -0.97 | **0.03** | -0.62 | **0.00** | 0.33 | 0.51 |
| **facet number** | -10.02 | 0.44 | 0.30 | 0.71 | **0.00** | -0.16 | 0.68 |
| **mean eye parameter (µm.rad)** | 23.16 | -1.28 | **0.03** | -0.28 | 0.26 | 0.31 | 0.68 |

**Table S7: detailed results of presence in forest on visual traits in workers, males and queens.** Simple or multiple Bayesian linear regressions including presence in forest and/or eye size were modelled on each visual trait. The lowest Deviance Information Criterion (*DIC*) indicates the best model for each trait (highlighted in beige). Significant positive or negative effects are reported in blue or red (respectively). The significance was based on the Bayesian p-value ($p_{MCMC}$ < 0.05). The phylogenetic heritability was the proportion of the posterior covariance explained by phylogeny.

Effects of *presence in forest*

| WORKERS | **DIC** | **Slope** | **$p_{MCMC}$** | **Phylogenetic heritability (%)** |
|---|---|---|---|---|
| **mean facet diameter (μm)** | 91.40 | 0.56 | 0.14 | 0.49 |
| **mean IO angle (°)** | 59.19 | 0.61 | 0.16 | 0.43 |
| **facet number** | 83.70 | -0.16 | 0.69 | 0.32 |
| **mean eye parameter (μm.rad)** | 88.87 | 1.20 | **0.00** | 0.23 |

| MALES | **DIC** | **Slope** | **$p_{MCMC}$** | **Phylogenetic heritability (%)** |
|---|---|---|---|---|
| **mean facet diameter (μm)** | 29.09 | -0.01 | 0.98 | 0.41 |
| **mean IO angle (°)** | -11.10 | 0.38 | 0.51 | 0.51 |
| **facet number** | 21.39 | -0.73 | 0.16 | 0.44 |
| **mean eye parameter (μm.rad)** | 22.41 | 0.38 | 0.51 | 0.36 |

| QUEENS | **DIC** | **Slope** | **$p_{MCMC}$** | **Phylogenetic heritability (%)** |
|---|---|---|---|---|
| **mean facet diameter (μm)** | -0.38 | -0.68 | 0.35 | 0.54 |
| **mean IO angle (°)** | -2.04 | 0.96 | 0.19 | 0.46 |
| **facet number** | 0.32 | 0.21 | 0.79 | 0.47 |
| **mean eye parameter (μm.rad)** | 0.23 | 0.79 | 0.29 | 0.54 |

Effects of eye size

| WORKERS | **DIC** | **Slope** | **$p_{MCMC}$** | **Phylogenetic heritability (%)** |
|---|---|---|---|---|
| **mean facet diameter (μm)** | 14.28 | 0.98 | **0.00** | 1.17 |
| **mean IO angle (°)** | 41.98 | -0.46 | **0.00** | 0.44 |
| **facet number** | 16.59 | 0.90 | **0.00** | 1.04 |
| **mean eye parameter (μm.rad)** | 69.29 | 0.62 | **0.00** | 0.40 |

| MALES | **DIC** | **Slope** | **$p_{MCMC}$** | **Phylogenetic heritability (%)** |
|---|---|---|---|---|
| **mean facet diameter (μm)** | 19.05 | 0.81 | **0.00** | 0.53 |
| **mean IO angle (°)** | -27.42 | -0.59 | **0.02** | 0.57 |
| **facet number** | 4.38 | 0.90 | **0.00** | 0.79 |
| **mean eye parameter (μm.rad)** | 19.12 | -0.17 | 0.52 | 0.36 |

| QUEENS | **DIC** | **Slope** | **$p_{MCMC}$** | **Phylogenetic heritability (%)** |
|---|---|---|---|---|
| **mean facet diameter (μm)** | -5.86 | 0.78 | **0.00** | 0.56 |
| **mean IO angle (°)** | -5.10 | -0.76 | **0.01** | 0.55 |
| **facet number** | -5.36 | 0.83 | **0.00** | 0.55 |
| **mean eye parameter (μm.rad)** | -1.35 | -0.39 | 0.25 | 0.51 |

Effects of *presence in forest* and eye size

| WORKERS | **DIC** | ***presence in forest* slope** | **$p_{MCMC}$** | **eye size slope** | **$p_{MCMC}$** |
|---|---|---|---|---|---|
| **mean facet diameter (μm)** | 12.23 | 0.32 | **0.02** | 0.95 | **0.00** |
| **mean IO angle (°)** | 41.85 | 0.82 | **0.03** | -0.51 | **0.00** |

| | | | | | |
|---|---|---|---|---|---|
| **facet number** | 14.43 | -0.38 | **0.01** | 0.92 | **0.00** |
| **mean eye parameter (µm.rad)** | 69.27 | 1.10 | **0.01** | 0.53 | **0.00** |

| MALES | **DIC** | ***presence in forest* slope** | **$p_{MCMC}$** | **eye size slope** | **$p_{MCMC}$** |
|---|---|---|---|---|---|
| **mean facet diameter (µm)** | 20.70 | 0.41 | 0.20 | 0.86 | **0.00** |
| **mean IO angle (°)** | -29.16 | 0.12 | 0.80 | -0.59 | **0.02** |
| **facet number** | 6.30 | -0.30 | 0.21 | 0.86 | 0.00 |
| **mean eye parameter (µm.rad)** | 21.53 | 0.31 | 0.61 | -0.14 | 0.62 |

| QUEENS | **DIC** | ***presence in forest* slope** | **$p_{MCMC}$** | **eye size slope** | **$p_{MCMC}$** |
|---|---|---|---|---|---|
| **mean facet diameter (µm)** | -6.21 | -0.49 | 0.28 | 0.76 | **0.01** |
| **mean IO angle (°)** | -3.55 | 0.79 | 0.10 | -0.72 | **0.01** |
| **facet number** | -5.12 | 0.44 | 0.37 | 0.86 | **0.01** |
| **mean eye parameter (µm.rad)** | 0.76 | 0.70 | 0.35 | -0.36 | 0.28 |

Effects of *presence in forest* and eye size and of their interaction

| WORKERS | **DIC** | ***presence in forest* slope** | **$p_{MCMC}$** | **eye size slope** | **$p_{MCMC}$** | **Interaction *presence in forest* and eye size slope** | **$p_{MCMC}$** |
|---|---|---|---|---|---|---|---|
| **mean facet diameter (µm)** | 3.93 | 0.25 | 0.08 | 1.15 | **0.00** | -0.29 | **0.02** |
| **mean IO angle (°)** | 45.23 | 0.79 | **0.02** | -0.46 | **0.01** | -0.11 | 0.64 |
| **facet number** | 9.17 | -0.33 | **0.04** | 0.76 | **0.00** | 0.24 | 0.07 |
| **mean eye parameter (µm.rad)** | 67.58 | 0.97 | **0.02** | 0.83 | **0.00** | -0.50 | 0.08 |

| MALES | DIC | *presence in forest* slope | $p_{MCMC}$ | eye size slope | $p_{MCMC}$ | Interaction *presence in forest* and eye size slope | $p_{MCMC}$ |
|---|---|---|---|---|---|---|---|
| **mean facet diameter ($\mu$m)** | 20.88 | 0.45 | 0.18 | 0.98 | **0.01** | -0.19 | 0.59 |
| **mean IO angle (°)** | -34.81 | 0.24 | 0.67 | -0.24 | 0.58 | -0.49 | 0.37 |
| **facet number** | 6.39 | -0.32 | 0.20 | 0.79 | **0.00** | 0.11 | 0.65 |
| **mean eye parameter ($\mu$m.rad)** | 20.16 | 0.45 | 0.48 | 0.23 | 0.67 | -0.52 | 0.41 |

| QUEENS | DIC | *presence in forest* slope | $p_{MCMC}$ | eye size slope | $p_{MCMC}$ | Interaction *presence in forest* and eye size slope | $p_{MCMC}$ |
|---|---|---|---|---|---|---|---|
| **mean facet diameter ($\mu$m)** | -5.34 | -0.59 | 0.20 | 0.27 | 0.53 | 0.60 | 0.25 |
| **mean IO angle (°)** | -3.75 | 0.85 | 0.10 | -0.39 | 0.41 | -0.41 | 0.44 |
| **facet number** | -4.71 | 0.54 | 0.29 | 1.31 | **0.03** | -0.57 | 0.30 |
| **mean eye parameter ($\mu$m.rad)** | 1.79 | 0.71 | 0.38 | -0.32 | 0.70 | -0.05 | 0.94 |

**Table S8: detailed results of presence in forest and eye size on visual traits in all workers except *B. transversalis*.** Multiple Bayesian linear regressions including presence in forest and eye size were modelled on each visual trait. Significant positive or negative effects are reported in blue or red (respectively). The significance was based on the Bayesian p-value ($p_{MCMC}$ < 0.05).

| WORKERS (without *B. transversalis*) | **DIC** | ***presence in forest* slope** | **$p_{MCMC}$** | **eye size slope** | **$p_{MCMC}$** | **Interaction *presence in forest* and eye size slope** | **$p_{MCMC}$** |
|---|---|---|---|---|---|---|---|
| **mean facet diameter ($\mu$m)** | 28.90 | 0.45 | **0.03** | 1.06 | **0.00** | -0.31 | **0.03** |
| **mean IO angle (º)** | 50.70 | 0.88 | **0.02** | -0.32 | **0.02** | -0.07 | 0.76 |
| **facet number** | 30.12 | -0.51 | **0.01** | 0.67 | **0.00** | 0.27 | 0.06 |
| **mean eye parameter ($\mu$m.rad)** | 66.44 | 1.13 | **0.01** | 0.55 | **0.01** | -0.31 | |

**Table S9: detailed results of the forest affinity index on visual traits in workers.** Simple or multiple Bayesian linear regressions including presence in forest and/or eye size were modelled on each visual trait. The lowest Deviance Information Criterion (*DIC*) indicates the best model for each trait (highlighted in beige). Significant positive or negative effects are reported in blue or red (respectively). The significance was based on the Bayesian p-value ($p_{MCMC} < 0.05$). The phylogenetic heritability was the proportion of the posterior covariance explained by phylogeny.

Effects of *forest affinity index*

| WORKERS | **DIC** | **Slope** | $p_{MCMC}$ | **Phylogenetic heritability (%)** |
|---|---|---|---|---|
| **mean facet diameter ($\mu$m)** | 86.56 | 0.53 | 0.00 | 0.31 |
| **mean IO angle (º)** | 59.76 | 0.00 | 1.00 | 0.33 |
| **facet number** | 83.72 | 0.29 | 0.13 | 0.44 |
| **mean eye parameter ($\mu$m.rad)** | 90.85 | 0.45 | 0.01 | 0.4 |

Effects of eye size

| WORKERS | **DIC** | **Slope** | $p_{MCMC}$ | **Phylogenetic heritability (%)** |
|---|---|---|---|---|
| **mean facet diameter ($\mu$m)** | 14.28 | 0.98 | **0.00** | 1.17 |
| **mean IO angle (º)** | 41.98 | -0.46 | **0.00** | 0.44 |
| **facet number** | 16.59 | 0.90 | **0.00** | 1.04 |
| **mean eye parameter ($\mu$m.rad)** | 69.29 | 0.62 | **0.00** | 0.40 |

Effects of *forest affinity index* and eye size

| WORKERS | **DIC** | ***forest affinity index* slope** | $p_{MCMC}$ | **eye size slope** | $p_{MCMC}$ |
|---|---|---|---|---|---|
| **mean facet diameter ($\mu$m)** | 14.67 | 0.09 | 0.23 | 0.94 | **0.00** |
| **mean IO angle (º)** | 42.93 | 0.20 | 0.26 | -0.50 | **0.00** |
| **facet number** | 16.45 | -0.13 | 0.12 | 0.95 | **0.00** |
| **mean eye parameter ($\mu$m.rad)** | 70.50 | 0.25 | 0.28 | 0.55 | **0.01** |

Effects of *forest affinity index* and eye size and of their interaction

| WORKERS | **DIC** | ***forest affinity index* slope** | $p_{MCMC}$ | **eye size slope** | $p_{MCMC}$ | **Interaction *forest affinity index* and eye size slope** | $p_{MCMC}$ |
|---|---|---|---|---|---|---|---|
| **mean facet diameter ($\mu$m)** | 14.00 | 0.15 | 0.08 | 0.97 | **0.00** | -0.08 | 0.13 |
| **mean IO angle (º)** | 39.27 | 0.39 | 0.06 | -0.52 | **0.00** | -0.20 | 0.07 |
| **facet number** | 16.34 | -0.19 | 0.05 | 0.93 | **0.00** | 0.07 | 0.22 |
| **mean eye parameter ($\mu$m.rad)** | 64.59 | 0.53 | **0.04** | 0.57 | **0.00** | -0.31 | **0.03** |
